# Phosphatidylserine receptors are critical for lethal infection by Crimean-Congo hemorrhagic fever virus

**DOI:** 10.64898/2026.10.29.762781

**Authors:** Jacob Berrigan, Stephanie R. Monticelli, Matt Daileg Ramirez, Paige T. Donovan, Taylor Powell, Ana I. Kuehne, Russell R. Bakken, Sarah B. Sulkosky, Xiankun Zeng, Christopher P. Stefan, Susan R. Coyne, Andrew S. Herbert, Wendy J. Maury, Kartik Chandran, Ezgi Kasikci

## Abstract

The tick-borne Crimean-Congo hemorrhagic fever virus (CCHFV) causes severe and often fatal hemorrhagic disease for which no approved countermeasures are available. Herein, we identify the phosphatidylserine receptors (PSRs) AXL, TIM-1, and TIM-4 as critical entry host factors for CCHFV in cell lines and liver-derived primary cell cultures. Small-molecule and antibody-based PSR inhibitors greatly reduced viral infection in cell culture. Germline knockout of individual PSR genes afforded partial protection against CCHFV infection, disease, and death in a murine model of lethal viral challenge. Mice engineered to lack two PSRs were completely protected from challenge, with viremia and viral replication, pathology, and inflammation in the liver and spleen reduced to baseline. Therefore, PSRs are collectively indispensable for CCHFV infection and pathogenesis. We conclude that PSRs are not just cell entry factors but *bona fide* determinants of infection, dissemination, and disease by CCHFV and legitimate targets for the development of new antiviral countermeasures.

## Introduction

Crimean-Congo hemorrhagic fever virus (CCHFV) belongs to the genus *Orthonairovirus* in the family *Nairoviridae* of enveloped, segmented, negative-strand RNA viruses (hereafter, orthonairoviruses). CCHFV is primarily spread by hyalommid ticks and can infect a wide variety of mammals and birds[1]. Its broad geographic distribution reflects those of its arthropod vectors and vertebrate hosts, with a range extending from sub-Saharan Africa to northeastern China[2–4]. CCHFV is a high-consequence pathogen associated with lethal disease in humans with case-fatality rates approaching 40%. Human cases are characterized by febrile illness, thrombocytopenia and multi-organ failure[5]. There are currently no specific FDA- or EMA-approved countermeasures for CCHFV; however, multiple vaccines and monoclonal antibody treatments are currently in development[6].

The tri-segmented negative-strand CCHFV genome (Genbank Accession: NC_005302 [S], NC_005300 [M] and NC_005301 [L]) consists of the S segment encoding the nucleoprotein, the M segment encoding the glycoprotein complex (GPC), and the L segment encoding the RNA-dependent RNA polymerase. The GPC precursor polypeptide is co- and post-translationally cleaved by cellular proteases, including signal peptidase and the proprotein convertases subtilisin/kexin isozyme-1 (SKI-1) and furin, to yield the mature glycoprotein complex comprised of three subunits: GP38, Gn, and Gc[7–9]. Recently solved structures of soluble forms of a GP38/Gn/Gc complex and a post-fusion Gc homotrimer have provided insights into the biogenesis, architecture, and entry-related deployment of the structural unit that likely constitutes the (currently undefined) GPC lattice at the surface of viral particles[10–12]. GP38 also exists in a “secreted” (non-virion incorporated) form whose biogenesis is unclear and was recently shown to induce vascular hyperpermeability (‘leak’) *in vitro* and *in vivo*, suggesting that it is a key player in viral dissemination and/or pathogenesis[13].

Considerable recent efforts have aimed at uncovering the cellular molecules that interact with viral particles to mediate CCHFV entry and influence infection and pathogenesis *in vivo.* Three independent studies identified the low-density lipoprotein receptor (LDLR) as an entry factor for CCHFV. Two of these studies [14, 15] suggested that LDLR may bind directly to Gc, whereas the third [16] did not find evidence for a direct GPC:LDLR interaction but proposed instead that virion membrane-associated apolipoprotein E (APOE) bridges viral particles and LDLR. Xu and Monteil showed that germline *Ldlr* loss affords partial protection in an immunodeficient murine model of lethal CCHFV challenge[14, 15]. Ma and co-workers identified lactadherin (milk fat globule-EGF factor 8 protein; MFGE8), a secreted glycoprotein that bridges phosphatidylserine and integrins, as a candidate CCHFV entry factor, but they did not evaluate its role *in vivo*[17]. Taken together, therefore, the precise mechanistic roles of these host factors in CCHFV cell entry remain incompletely understood, as are their implications for viral virulence and pathogenesis. Moreover, other CCHFV host factors likely await discovery.

To identify additional human genes required for orthonairovirus entry, we performed genome-wide CRISPR/Cas9 knockout screens with a replication-competent recombinant vesicular stomatitis virus bearing the CCHFV-Ibar10200 GPC (rVSV-CCHFV GPC) and Hazara Virus (HAZV), a related avirulent orthonairovirus[18]. These screens identified the TAM receptor AXL as an orthonairovirus entry factor. We show herein that AXL, T-cell immunoglobulin and mucin domain containing protein-1 (TIM-1), a structurally unrelated phosphatidylserine receptor (PSR), and the TIM-1 paralog TIM-4 mediate CCHFV entry into immortalized cell lines and primary human liver cells through interactions with phosphatidylserine (PtdSer) in the viral envelope. Although roles for AXL, TIM-1, TIM-4, and other PSRs in cell entry have also been described for other enveloped viruses *in cellulo*, relatively limited evidence exists that cellular recognition of viral envelope PtdSer is a major mechanism for viral infection and pathogenesis *in vivo*[19–24]. Crucially, therefore, we present evidence that AXL, TIM-1, and TIM-4 are collectively indispensable for CCHFV infection and pathogenesis in a murine model of lethal challenge.

## Results

### Genome-wide CRISPR/Cas9 screens identify the receptor tyrosine kinase and PSR AXL as a candidate host factor for orthonairovirus infection

We used the Brunello CRISPR/Cas9 knockout library to screen U2OS human osteosarcoma cells and SW-13 human adrenocortical carcinoma cells for host genes whose loss afforded cell survival after infection with a replication-competent vesicular stomatitis virus bearing the CCHFV IbAr10200 glycoprotein complex (rVSV-CCHFV GPC [IbAr]) and HAZV, respectively[18, 25–28]. Overlapping sets of genes were enriched in the two virus-resistant populations (Fig. 1a-b and Data file S1). The products of these genes could be grouped into distinct functional categories/pathways, including components of the ER membrane complex (EMC); the Guided Entry of Tail-anchored proteins (GET) pathway for ER membrane insertion of tail-anchored membrane proteins; proteins involved in endo/lysosomal biogenesis, acidification, and trafficking; and components of the Type I interferon signaling pathway (Fig. 1c). The gene encoding AXL, a receptor tyrosine kinase and PSR, was the single most enriched cell-surface hit in both rVSV-CCHFV GPC and HAZV screens and was selected for further study.

**Figure 1.**
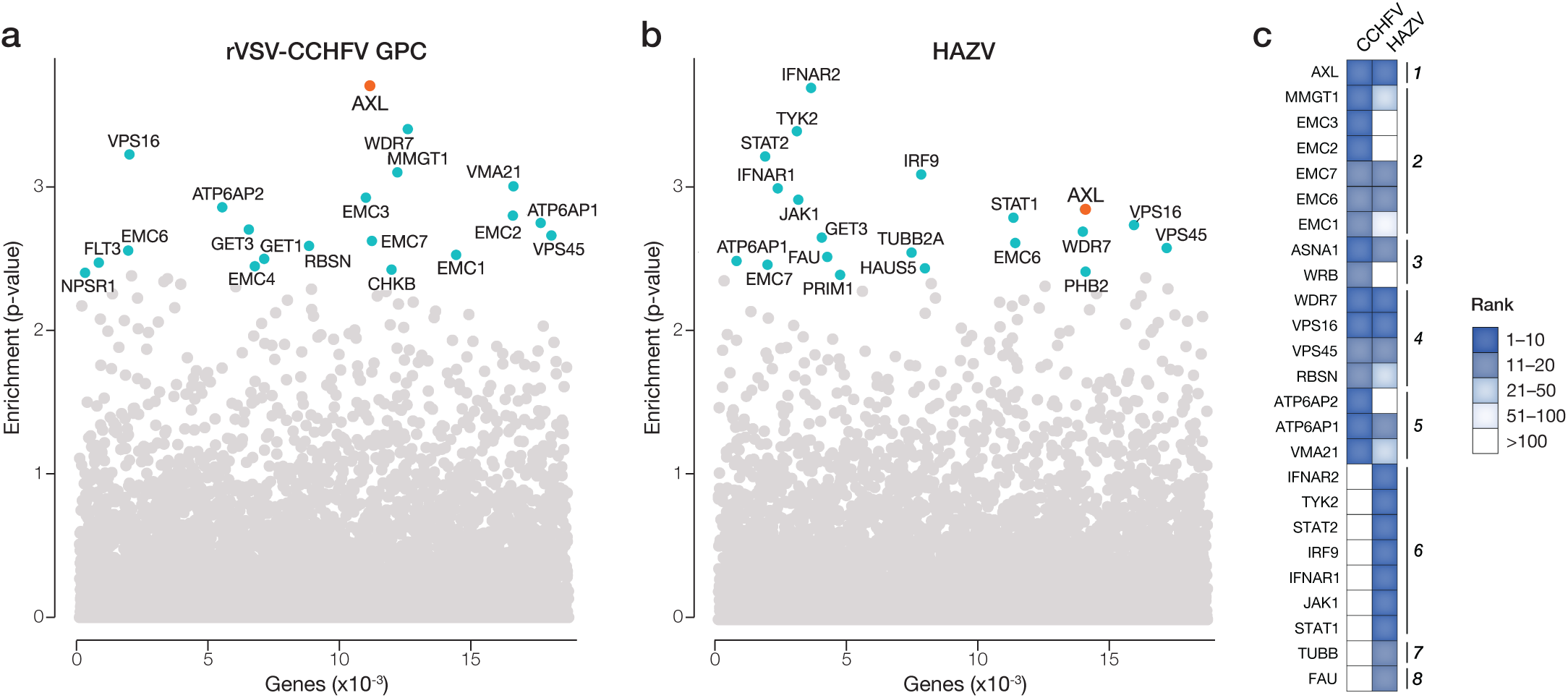
Genome-wide loss-of-function CRISPR/Cas9 screens identify AXL as a candidate factor for rVSV-CCHFV GPC and HAZV infection. (a-b) CRISPR/Cas9 screens with rVSV-CCHFV (IbAr10200) in human U2OS cells (a) and with HAZV in human SW-13 cells (b). Hit significance was determined with MAGeCK [27]. Genes with enrichment p-values >4 in the GeneRaMeN package [85] are colored and grouped by pathway. Combined results from two biological replicates are shown. (c) Heat map illustrating the top 20 hits from the rVSV-CCHFV screen ranked by enrichment score. The corresponding genes from the HAZV screen are also shown with their enrichment scores.

### Knockout of PSRs in immortalized cell lines reduces CCHFV entry and infection

AXL has been previously implicated in cell entry by distinct enveloped viruses by virtue of its capacity to recognize Gas6, a PtdSer-binding adapter that can recruit virions, driving AXL-mediated viral attachment and internalization into cells[29, 30]. To investigate the role of AXL in CCHFV entry, we generated clonal *AXL*-knockout (KO) cell lines using CRISPR/Cas9 engineering (Table S1). We selected A549 and Vero cells for our initial studies because both cell lines express AXL at high levels[31]. Biallellic *AXL* KO was confirmed by long-read sequencing of PCR amplicons spanning the single-guide RNA (sgRNA) target sites, western blot, and immunofluorescence staining (Figs. S1-S2). Control (empty vector; EV) and *AXL*-KO A549 cell lines were then exposed to rVSV-CCHFV GPC (IbAr). We observed a substantial reduction in infection in the KO cells (Fig. 2a). This infectivity loss was AXL-specific, because it could be largely restored in KO cells ectopically expressing AXL (complemented; CP) (Fig. 2a). These findings extended to transcription-and-entry competent virus-like particles (tecVLPs) that have been widely used in the field as CCHFV surrogates [32, 33](Fig. 2b). Most important, *AXL* KO and complementation respectively reduced and restored infection by authentic CCHFV (IbAr) (Fig. 2c). Similar results were observed in Vero cells upon *AXL* KO (Fig. S3a).

**Figure 2.**
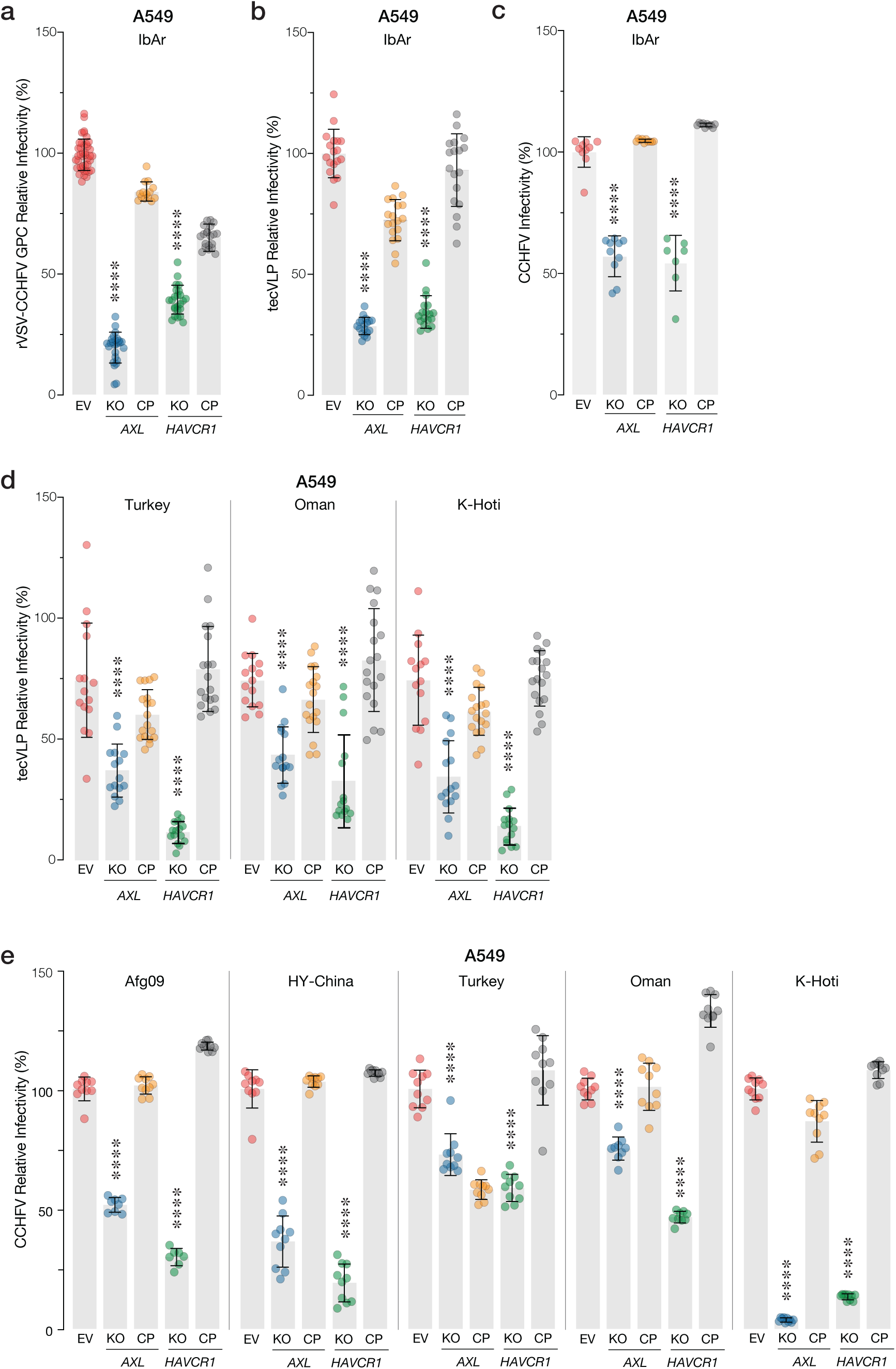
Effects of AXL and TIM-1 loss on CCHFV entry and infection. Infection of A549 cell clones knocked out (KO) for *AXL* or *HAVCR1* (encoding TIM-1). **(a)** Cells were exposed to rVSV-CCHFV GPC (MOI, 0.2 IU/cell) for 1 h prior to addition of NH4Cl to stop infection. Infected cells were enumerated at 16 hpi by immunofluorescence microscopy. **(b, d)** Cells were exposed to equal volumes of TCID,。-normalized CCHFV tecVLPs, and transduction was measured at 16 hpi by luminometry. **(c, e)** Cells were exposed to authentic CCHFV (MOI, 0.2 PFU/cell), and infected cells were enumerated by immunofluorescence microscopy. Means ± s.d. from three independent experiments are shown. Groups (WT vs. others) were compared by one-way ANOVA with Dunnett’s post hoc test.

To examine the viral strain-dependent behavior of CCHFV in these cell lines, we next exposed them to tecVLPs bearing GPC proteins from strains belonging to different viral clades. Cell entry mediated by the divergent GPC proteins was reduced in the *AXL*-KO lines and could be restored upon AXL complementation (Fig. 2d and Fig. S3). Finally, although the quantitative reduction in infection varied by strain, we observed largely concordant results with divergent authentic CCHFVs (Fig. 2e).

TIM-1, a PSR structurally unrelated to AXL, has also been implicated in viral entry, and *HAVCR1*, the gene encoding TIM-1, was a hit in a recently published CRISPR screen for CCHFV host factors[17]. However, the role of TIM-1 was not evaluated in that study. Accordingly, we generated clonal *HAVCR1* KO cell lines and examined their susceptibility to CCHFV entry and infection, as described for *AXL* above. The effects of TIM-1 loss and complementation in rVSV-, tecVLP-, and authentic virus-based infection assays closely paralleled that seen for AXL (Fig. 2 and Figs. S1-S3). Together, these findings point to a broad role for the PSRs AXL and TIM-1 in CCHFV entry and infection in immortalized cell lines.

### Overexpression of AXL and TIM-1 in immortalized cell lines boosts CCHFV entry and infection

SW-13 cells endogenously express low levels of AXL and TIM-1 (Fig. S3). To further corroborate the roles of AXL and TIM-1 as CCHFV entry factors, we assessed the effects of their over-expression in SW-13 cells (Fig. S3) on CCHFV entry and infection. We noted significant increases in IbAr GPC-dependent entry and/or infection in the rVSV-, tecVLP- and authentic virus-based assays upon over-expression of AXL or TIM-1, although the degree of the boost varied somewhat in each system (Fig. 3a-c). Similar, if variable, AXL- and TIM-1-dependent infection boosts were observed with divergent authentic CCHFVs (Fig. 3c). Whether the strain-dependent differences in these assays reflect true biological differences in host factor use or simply differences in the compositions of specific viral preparations remains unclear at present (see Discussion). By contrast, over-expression of Gas6, a PtdSer-binding adapter for AXL, had no effect, possibly because Gas6 was not limiting in this assay system. Importantly, over-expression of a TIM-1 mutant deficient in PtdSer-binding (N115D/N116D) did not enhance rVSV-CCHFV GPC entry (Fig. 3a)[20, 34], strongly suggesting that the entry-enhancing effect of TIM-1 over-expression requires its interaction with viral membrane PtdSer (also see Fig. 6 below).

**Figure 3.**
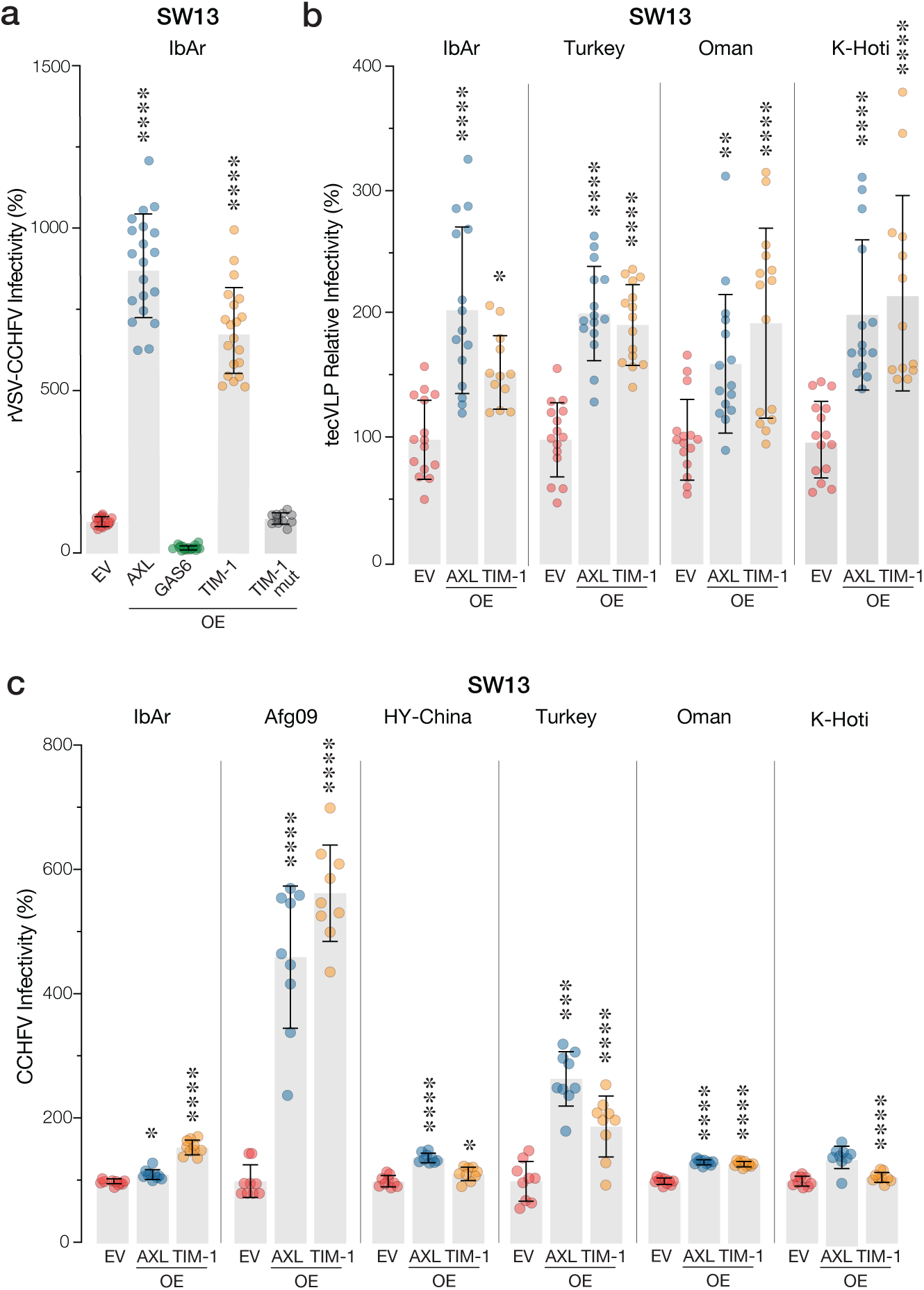
Effects of AXL and TIM-1 overexpression on CCHFV infection. Infection of SW-13 cells overexpressing AXL or TIM-1. **(a)** cells were exposed to rVSV-CCHFV GPC (MOI, 0.2 IU/cell for 1 h prior to addition of NH_4_Cl to stop infection). Infected cells were enumerated at 16 hpi by immuno-fluorescence microscopy. (**b**), Cells were exposed to equal volumes of TCID,。-normalized CCHFV tecVLPs, and transduction was measured at 16 hpi by luminometry. **(c)** Cells were exposed to authentic CCHFV (MOI, 0.2 PFU/cell), and infected cells were enumerated by immunofluorescence microscopy. Means ± s.d. from three inde-pendent experiments are shown. Groups (WT vs. others) were compared by one-way ANOVA with Dunnett’s post hoc test.

### Depletion of AXL and TIM-1 in physiologically relevant cell types reduces CCHFV entry

Work in murine and nonhuman primate models indicates that hepatocytes, endothelial cells, and Kupffer cells are major targets of CCHFV infection in the liver[35–39]. To begin to define the roles of AXL and TIM-1 for viral entry in primary cell types, we used lentiviral vectors encoding short-hairpin RNAs (shRNAs) to stably knock down mRNAs expressing these genes. First, we confirmed the effectiveness of the shRNA constructs in reducing protein expression and viral infection in A549 cells (Fig. S4a-b). We then used these vectors to knock down expression of AXL and TIM-1 in human umbilical vein endothelial cells (HUVEC), a widely used primary human endothelial cell model (Fig. S4c), and then exposed them to rVSV-CCHFV GPC (IbAr)[40]. rVSV-CCHFV infection was greatly reduced in cells expressing shRNAs targeting AXL and TIM-1 relative to their counterparts expressing a non-targeting shRNA (scramble; SCR) (Fig. 4a). Having established conditions for functional depletion of AXL and TIM-1 in HUVEC, we next performed these knockdowns in pooled human primary liver endothelial cells (HLECP1) from five donors and challenged them with tecVLPs bearing IbAr10200 or Oman GPC. tecVLP entry was substantially but selectively reduced in the AXL- and TIM-1-depleted cells (Fig. 4b and Fig. S4d). Infection was also reduced in the HLECP1 AXL and TIM-1 knockdown cells upon infection with HAZV (Fig. S4e). Finally, we assessed the importance of AXL and TIM-1 for CCHFV entry in HepaRG, terminally differentiated hepatic cells derived from a human hepatic progenitor cell line that retain many characteristics of primary human hepatocytes[40, 41]. Both rVSV-CCHFV GPC infection and tecVLP entry were reduced in HepaRG cells depleted of AXL and TIM-1 as above (Fig. 4c and Fig. S4f). We conclude that the PSRs AXL and TIM-1 contribute to CCHFV entry in multiple liver cell types associated with CCHFV infection and pathogenesis *in vivo*.

**Figure 4.**
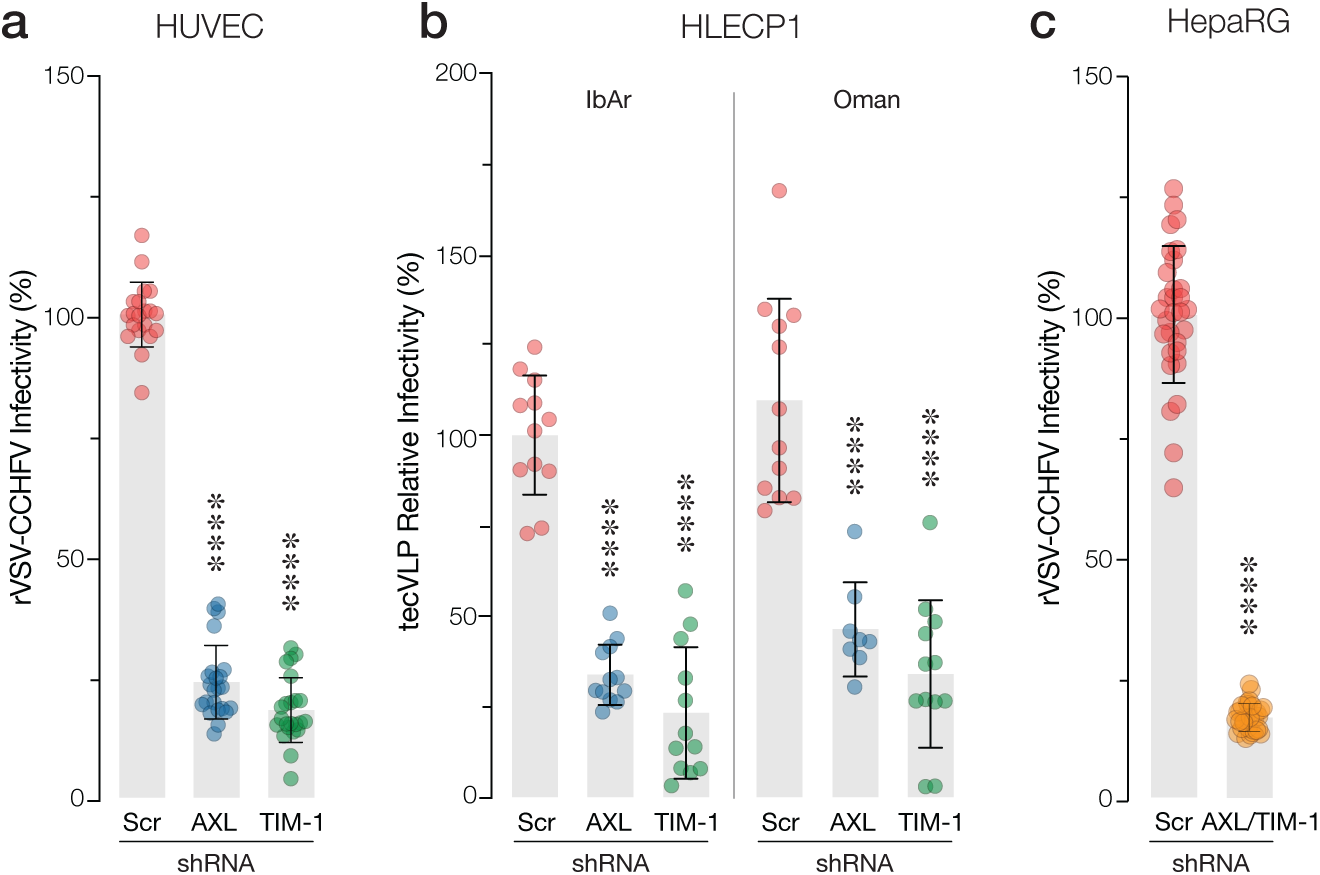
Knockdown of AXL and TIM-1 in physiologically relevant cell lines and primary cells. Infection of HUVEC, HLECP1, and HepaRG cells encoding short hairpin RNA (shRNA) constructs. HUVEC **(a)** and HepaRG **(b)** cells were exposed to rVSV-CCHFV GPC (MOI, 0.2 IU/cell) for 1 h prior to addition of NH_4_Cl to stop infection. Infected cells were enumerated at 16 hpi by immunofluorescence microscopy. **(b)** HLECP1 cells were exposed to equal volumes of TCID,。-normalized CCHFV tecVLPs, and transduction was measured at 16 hpi by luminometry. Means ± s.d. from three independent experiments are shown. Groups (WT vs. others) were compared by one-way ANOVA with Dunnett’s post hoc test.

### Cell entry by CCHFV is dependent on PtdSer binding and AXL-dependent entry requires its tyrosine kinase activity

We postulated that AXL and TIM-1 drive CCHFV entry by recognizing PtdSer in the outer leaflet of the virion membrane, thereby mediating viral attachment and internalization into cells, as has been established for many enveloped viruses that can coöpt these PSRs[29, 30, 34, 42].

Indeed, our finding that a TIM-1 mutant deficient in PtdSer binding could not boost CCHFV entry (Fig. 3a) provided initial support for this mechanism. To further evaluate the role of PtdSer, we sought to out-compete PSR:virion interactions by saturating cell-surface PSRs with excess PtdSer. Specifically, we pre-incubated Vero cells with synthetic liposomes comprising PtdSer, as described previously[24, 43], and then exposed them to rVSV-CCHFV GPC (IbAr). Liposomes constituted with phosphatidylcholine (PtdCho) were used as a control. PtdSer liposomes blocked CCHFV entry in a dose-dependent manner, whereas the PtdCho liposomes had little effect (Fig. 5a). Divergent authentic CCHFVs were similarly sensitive to PtdSer liposomes but not PtdCho liposomes (Fig. 5b). These findings support the hypothesis that CCHFV coöpts the PtdSer-binding activity of AXL (through Gas6) and TIM-1 for entry.

**Figure 5.**
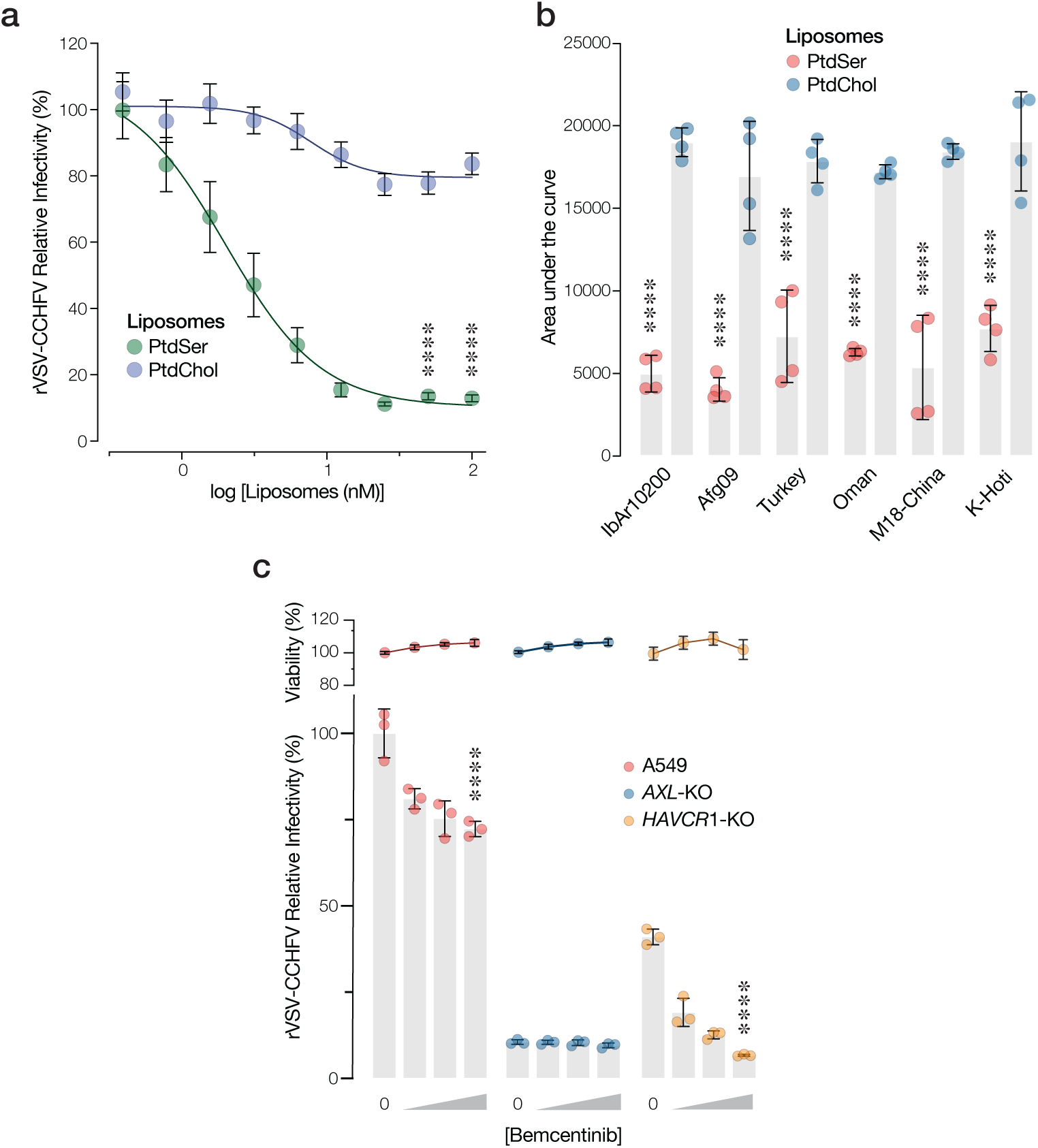
Mechanism of PSR-mediated CCHFV entry. (**a**) Vero cells were pretreated with increasing concentrations of PtdSer or PtdChol liposomes for 1 h prior to infection and then exposed to rVSV-CCHFV GPC (MOI, 0.2 IU/cell) for 1 h prior to addition of NH_4_Cl to stop infection. Infected cells were enumerated at 16 h post-infection (hpi) by immunofluorescence microscopy. Means ± s.d. from 3 independent experiments are shown. and groups were compared at the two highest mAb concentrations by unpaired Welch’s T-test. (**b**) Vero cells were pretreated with varying concentrations of PS or PC for 1 hour prior to infection with authentic CCHFV (MOI, 0.2 PFU/cell), and infected cells were enumerated by immunofluorescence microscopy. (**c**) A549 cells were pretreated with bemcentinib for 1 h and then exposed to rVSV-CCHFV GPC (MOI, 0.2 IU/cell) for 1 h prior to addition of NH_4_Cl to stop infection. Infected cells were enumerated at 16 hpi by immunofluorescence microscopy. Means ± s.d. from three independent experiments are shown. and groups (bemcentinib [0 vs. 2.5) were compared by one-way ANOVA with Dunnett’s post hoc test. Viability of cells was quantified post treatment and infection via DAPI signal from adherent cells relative to untreated and uninfected cells.

Previous work has shown that PtdSer-containing vesicles cluster Gas6, allowing it to recognize its low-affinity binding site in AXL[44, 45]. This in turn drives AXL activation and downstream signaling[46, 47]. Bemcentinib (R428), a potent and highly selective small-molecule AXL inhibitor currently under clinical investigation, blocks AXL’s receptor tyrosine kinase activity, thereby preventing its autophosphorylation and activation in response to Gas6/PtdSer binding[48]. To test if CCHFV entry requires AXL activation, we assessed the effect of bemcentinib treatment on rVSV-CCHFV GPC infection. First, we identified a concentration range of bemcentinib that did not significantly reduce cell viability (Fig. 5c). Next, we pre-treated A549 WT, *AXL*-KO, and *HAVCR1*-KO cells under these conditions and then exposed them to rVSV-CCHFV GPC (Fig. 5c). Infection was only modestly inhibited in WT cells and not at all in *AXL*-KO cells. By contrast, we observed a substantial (≈90%) reduction in infection in *HAVCR1*-KO cells. Together, these results provide evidence that: (i) the activity of bemcentinib under these conditions is AXL-specific (lack of effect in *AXL*-KO cells); (ii) AXL-dependent CCHFV entry requires AXL activation through its receptor tyrosine kinase activity (strong effect in *HAVCR1*-KO cells); and (iii) AXL and TIM-1 play redundant roles in CCHFV entry (strong effect in *HAVCR1*-KO cells but modest effect in WT cells expressing both AXL and TIM-1).

### A cocktail of AXL- and TIM-1-specific human monoclonal antibodies blocks CCHFV GPC-dependent entry

We next evaluated the effects of AXL- and TIM-1–blocking monoclonal antibodies on CCHFV entry (Fig. 6a). Preincubation of Vero cells with increasing concentrations of either mAb alone reduced rVSV-CCHFV GPC infection modestly or not at all. In striking contrast, however, the combination of α-AXL and α-TIM-1 mAbs potently inhibited rVSV-CCHFV GPC infection in a dose-dependent manner (Fig. 46).

**Figure 6.**
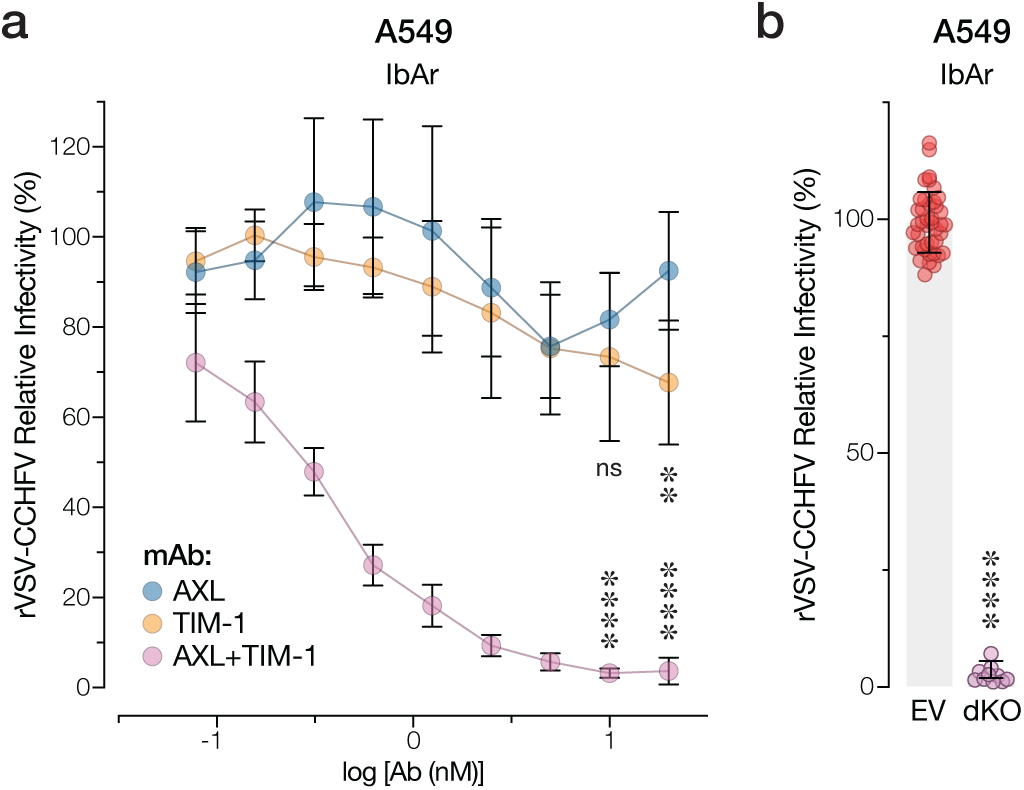
AXL- and TIM-1-specific mAbs block CCHFV entry. **(a)** Vero cells were pretreated with ɑ-AXL and/or ɑ-TIM-1 mAbs for 1 h and then exposed to rVSV-CCHFV GPC (MOI, 0.2 IU/cell) for 1 h prior to addition of NH_4_Cl to stop infection. Infected cells were enumerated at 16 hpi by immunofluorescence microscopy. Means ± s.d. from 3 independent experiments are shown. and groups were compared at the two highest mAb concentrations by two-way ANOVA with Tukey’s post hoc test. **(b)** A549 cells were exposed to rVSV-CCHFV GPC (MOI, 0.2 IU/cell) for 1 h prior to addition of NH_4_Cl to stop infection. Infected cells were enumerated at 16 hpi by immunofluorescence microscopy. Means ± s.d. from 3 independent experiments are shown. and groups were compared.

To complement these findings with a genetic approach, we engineered the KO of *HAVCR1* in a A549 *AXL*-KO background (Fig. 2). rVSV-CCHFV GPC infection was reduced by >90% in these *AXL/HAVCR1*-KO cells (Fig. 6b), whereas only ≍50–75% reductions were observed in the individual KO clones (see Fig. 2a). Together, these findings provide evidence that cell-surface AXL and TIM-1 play direct and overlapping roles in CCHFV entry and that mAbs simultaneously targeting both proteins can protect cells from infection.

### PSRs are critical for survival and disease progression in mice

Finally, we assessed the importance of PSRs for CCHFV virulence in germline double-knockout (dKO) mice. In addition to AXL and TIM-1 we also incorporated TIM-4, a closely related PSR with previously described entry factor activity, into our challenge studies[19, 42, 49]. These mice lacked the gene encoding interferon α/β receptor (*Ifnar1*; to render the mice susceptible to CCHFV) and an individual PSR-encoding gene: *Axl*, *Havcr1*, or *Timd4* (Fig. S5; see Methods for details)[22, 50]. *Ifnar1/Axl*-dKO, *Ifnar1/Havcr1*-dKO, and *Ifnar1*/*Timd4*-dKO mice (n=15-16 per group) were challenged intraperitoneally (i.p.) with CCHFV (Turkey2004; 10 plaque-forming units [PFU]), and survival, weight loss, and clinical scores were monitored for 28 days (Fig. 7a-b and Fig. S5). Five mice per group were sacrificed on day 5, and serum and organs were harvested, analyzed for infectious viral load and viral RNA by plaque assay (Fig. 7c) and qRT-PCR (Fig. 7d), respectively, and subjected to histopathological analysis (Fig. 7e-f). Loss of any of the PSRs promoted survival, with the strongest effects observed upon loss of TIM-1 (*Havcr1*) or TIM-4 (*Timd4*) (Fig. 7b). The findings regarding weight loss and clinical scores were consistent with those from the survival study (Fig. S5a-b). As expected, infectious virus was detectable in serum, visceral organs (spleen, liver and kidney), and the brain in the *Ifnar1*-KO control mice on day 5. Loss of two PSR genes was associated with reduced viral titers in all sampled tissues but was especially notable in the liver and spleen (Fig. 7c,d). Reductions in viral genome loads varied by tissue and PSR KO but were generally smaller than those in infectious titer (Fig. 7d), presumably reflecting that the RT-qPCR assay samples a larger pool of viral material that includes non-infectious virus, and intact and fragmented viral RNA localized within and outside cells. To directly visualize infected and/or virus-producing cells, we next stained viral genomic RNA in tissue sections by *in situ* hybridization (RNA-ISH) (Fig. 7e-f). High levels of viral RNA-positive cells were observed in all tissues in the *Ifnar1*-KO control group and were reduced by PSR KO, although these reductions were less pronounced in the *Axl*-KO mice, as also seen in the kidney and liver infectivity and genome titers. In keeping with these findings, PSR KO ameliorated signs of pathology and inflammation in the liver and spleen (Fig. 7f).

**Figure 7.**
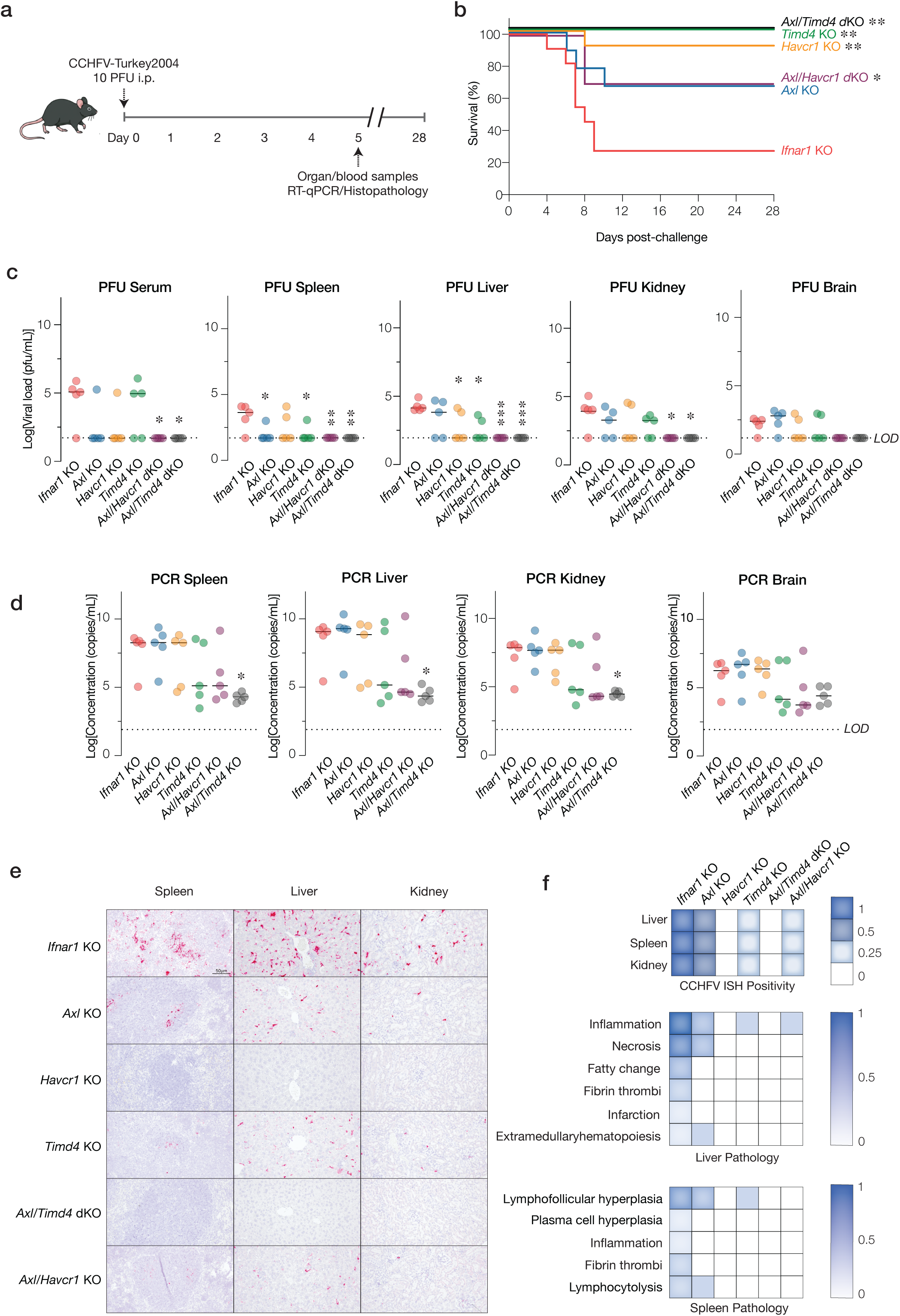
PSRs are crucial for CCHFV infection, disease, and lethality in mice. (**a**) *Ifnar1*-KO mice and PSR-KO mice generated in the *Ifnar1*-KO background (n=15-16 per strain) were challenged with Turkey2004 (10 PFU, i.p.). At 5 days post-challenge, 5 mice per group were euthanized and serum, liver, kidney, and spleen were harvested. (**b**) Mice were monitored for survival for 28 days. Survival was analyzed by two-sided Mantel-Cox (log-rank) tests. LOD, limit of detection. **(c)** Viral infectivity in the brain, kidney, spleen, liver and serum was quantified by plaque assay and compared with a Kruskal–Wallis test. **(d)** Viral genomes were quantified by RT-qPCR. **(e)** Viral RNA in in the spleen, liver, and kidney was detected by RNA-*in situ* hybridization (RNA-ISH). **(f)** Tissues were assessed for the incidence of RNA-ISH positivity and histologic lesions, scored as the proportion of affected animals per group (0 = none; 1 = all).

Given the additivity/redundancy among PSRs in mediating viral entry noted herein (Fig. 4) and elsewhere[51–53], we considered that PSR double-KO might accentuate the protective phenotypes observed in the PSR single-KO mice. Accordingly, we evaluated the virologic and pathologic trajectories of *Ifnar1/Axl/Havcr1* and *Ifnar1/Axl/Timd4* triple-KO (tKO) mice challenged with CCHFV as above. Both tKO lines were largely protected against challenge (Fig. 7b). Even more striking, *Axl/Timd4* loss completely prevented weight loss and clinical sings of disease (Fig. S5a-b). Further, PSR dKO reduced viral infectivity to baseline in serum and all sampled solid organs (Fig. 7d) and afforded greater diminutions in viral genome loads than the PSR single-KOs (Fig. 7d). This was especially remarkable in all the organs for *Axl/Timd4*-dKO and in serum for both *Axl/Timd4*-dKO and *Axl/Havcr1* dKO. In sum, our results establish the PSRs AXL, TIM-1, and TIM-4 as critical host factors for CCHFV virulence that drive viral infection, replication, and *in vivo* pathogenesis in an overlapping manner.

## Discussion

Herein, we investigated host requirements for CCHFV entry, infection, and virulence. Using comprehensive loss-of-function CRISPR-Cas9 screens and follow-up genetic studies, we show that the divergent PtdSer receptors (PSRs) AXL, TIM-1, and TIM-4 play overlapping roles in infection by the high-consequence orthonairovirus CCHFV. Our mechanistic studies indicate that the CCHFV entry activity of these PSRs requires their capacity to directly or indirectly bind PtdSer-containing cargo and the ligand-mediated intracellular signaling activity of AXL, consistent with a role for these receptors in viral attachment and internalization into cells (Figs. 2, 3). The CCHFV requirement for PSR activity was observed in multiple immortalized cell lines, primary endothelial cells of liver origin, and in terminally differentiated hepatic cells derived from a human liver progenitor cell line (Fig. 5). Further, it is generalizable to members of multiple divergent CCHFV clades and to the distinct orthonairovirus HAZV (Fig. S4). Most important, we provide biochemical evidence that antibodies targeting AXL and TIM-1 can block CCHFV entry *in cellulo* and genetic evidence that AXL, TIM-1, and TIM-4 are collectively indispensable for CCHFV replication and pathogenesis in a murine model of lethal CCHFV challenge (Fig. 7). This study is, to our knowledge, the first to evaluate the effects of single and double PSR gene KO on systemic replication and disease by an authentic hemorrhagic fever virus. We conclude that PSRs are not just cell entry factors but *bona fide* determinants of viral infection, dissemination, and disease and legitimate targets for the development of new antiviral countermeasures against CCHFV.

Previous work has identified multiple candidate host factors and receptors in CCHFV entry. Three groups independently uncovered the low-density lipoprotein receptor (LDLR) as a CCHFV entry factor, and two found a direct interaction between the CCHFV glycoprotein Gc and LDLR, whereas the third instead implicated an interaction between virion-bound apolipoprotein E (APOE) and LDLR[14–16]. More recently, Ma et al. identified MFGE8 (lactadherin), a secreted PtdSer-binding protein, as a CCHFV entry factor in immortalized cell lines but did not evaluate its requirement in primary cells or animal models[17].

What roles, if any, do these cell entry factors play in CCHFV infection and pathogenesis *in vivo*? Prior studies tested the requirement for LDLR in an immunosuppression-based murine model of lethal CCHFV challenge[14, 15]. However, only partial protection and partial reductions in viral replication in the serum, liver, and spleen in *Ldlr*-KO mice were observed, indicating the existence of other cell entry factors/pathways *in vivo*. Our findings herein with single PSR KOs in the *Ifnar1*-KO genetic background largely mirrored those in *Ldlr*-KO mice. Remarkably, however, double PSR-KO mice displayed an even greater degree of resistance to lethal viral infection, replication, and pathogenesis (Fig. 7). This was especially so in the *Axl/Timd4*-dKO mice, which were not only completely protected from challenge but also showed no detectable infectious virus in the serum, liver, and spleen, and greatly reduced viral RNA levels in these and other organs (Fig. 7). We conclude that PSR- and LDLR-mediated entry pathways are not truly redundant in the murine model for CCHFV. More work is needed to determine if they participate sequentially in CCHFV entry in the same cell types or instead independently mediate entry into distinct cell types in different tissues (also see below).

CCHFV can spread systemically in immunosuppressed murine models and can also replicate in multiple organ systems in humans, suggesting either that key cell entry factors are widely expressed, and/or that distinct suites of entry factors drive viral infection in specific cell types, possibly in a manner that changes over the course of disease [38, 54–56]. The broad tissue distribution of PSRs and our findings that they are required for CCHFV entry into multiple primary liver cell types (hepatocyte-like cells, liver endothelial cells), suggest that these molecules play ubiquitous roles in CCHFV infection *in vivo*. However, the distinct expression patterns of PSRs and LDLR in mice and humans also point to more complex scenarios. As a case in point, AXL and TIM-4 transcripts are highly expressed in splenic macrophages and liver-resident macrophages (Kupffer cells), whereas LDLR is more highly expressed in hepatocytes [57–60]. It is thus interesting to speculate that PSR-mediated infections of macrophages “seed” viral infection and amplify viral spread to other cell types and proinflammatory signaling in the liver and other organs. Intriguingly, recent work with HAZV in an immunocompromised murine model showed that Kupffer cells are crucial early targets of CCHFV infection and suggested that infection and IFNAR signaling in these cells synergize to drive viral spread in hepatocytes and liver injury[39]. Kupffer cells have also been suggested as important targets for infection and mediators of hepatic injury by Ebola virus, another enveloped virus that can use PSRs for infection (see below) [61]. Resolving these questions will require a more sophisticated understanding of the trajectory of viral dissemination, inflammation, and tissue injury during CCHFV infection.

Given the large body of research on PSRs in enveloped virus entry *in cellulo*, the evidence that these molecules influence viral virulence and pathogenesis is sparse and decidedly mixed. *Axl* loss partially protected mice from influenza A virus challenge and reduced lung pathology [23]. AXL was shown to play a role in Ebola virus (EBOV) infection in a human skin explant model, and *Axl* loss reduced replication by a surrogate recombinant vesicular stomatitis virus bearing EBOV GP (rVSV-EBOV GP) in a murine skin infection model, but its role in systemic infection was not evaluated [21, 62]. By contrast, despite abundant evidence for the TAM (TYRO3, AXL, MER) receptor tyrosine kinases as proviral entry factors for flaviviruses, arenaviruses, bunyaviruses, and filoviruses (among others) [29, 30, 63–66], studies with single or double TAM-KO mice have found these molecules to be largely dispensable [67–70]. To the contrary, TAM KOs appeared to worsen disease by Japanese encephalitis virus, West Nile virus, and La Crosse virus, potentially due to confounding effects related to the capacity of these molecules to modulate inflammation [71–73]. The *in vivo* evidence for TIM-1 is more promising: Brunton and co-workers found that *Havcr1*-KO afforded partial protection and reduced viral loads in multiple organs following challenge of mice with rVSV-Ebola virus GP [22]. However, challenge studies with authentic EBOV have not been reported. *Havcr1* loss also protected mice from lethal challenge by a neurotropic flavivirus, tick-borne encephalitis virus [74]. No *in vivo* studies investigating the role of TIM-4 have been reported to date. Based on our findings, we speculate that double-KO of a TAM and a “TIM” (TIM-1 or TIM-4) in mice might uncover significant proviral roles for these molecules in *in vivo* infection by enveloped viruses other than CCHFV.

Together with our findings *in vivo*, the potency of bemcentinib (Fig. 5c) and anti-PSR mAb cocktails (Fig. 6a) *in cellulo* raises the prospects for the development of PSR inhibitors as host-directed antivirals against CCHFV and other emerging orthonairoviruses. The clinical feasibility of targeting AXL for non-infectious disease indications is well established: multiple, specific small-molecule inhibitors, including bemcentinib, have been developed [48, 75], and a fully humanized anti-AXL mAb that blocks AXL:GAS6 binding, tilvestamab [76], has successfully completed Phase I clinical trials (NCT03795142). Together with prior work, our results suggest that pharmacological AXL blockade would likely act by two independent mechanisms: inhibition of cell entry and derepression of the Type I interferon response [77, 78]. Interestingly, most published studies have attributed the antiviral effects of AXL to the immune signaling axis alone [79–81], whereas our studies in *Ifnar1^-/-^* mice at least partially decouple the two mechanisms to demonstrate direct proviral roles for AXL, TIM-1, and TIM-4 (Fig. 7). Anti-TIM1 and anti-TIM4 mAbs are not as advanced, but TIM-1’s role in T-cell activation has been targeted by mAbs in proof-of-concept studies to enhance transplantation tolerance and ameliorate graft-vs.-host-disease [82, 83]. Two important considerations for the development of PSR-targeting antivirals for CCHFV are the likely need for simultaneous inhibition of both “TIM” and “TAM” arms as shown herein, and the challenges inherent in inhibiting host molecules with complex, context- and cell type-dependent roles in immune signaling and immune cell homeostasis [58, 59, 84, 85].

## Materials and Methods

### Cell culture

African green monkey kidney Vero cells [American Type Culture Collection (ATCC), Cat# CCL-81] and VeroE6 cells [ATCC, Cat#CRL-1586] were cultured in high-glucose Dulbecco’s Modified Eagle Medium (DMEM; Gibco, Cat# 11965-092) supplemented with 2% heat-inactivated fetal bovine serum (FBS; GeminiBio, Cat# 100-106, Lot A123000), and 1% penicillin-streptomycin (PS; Gibco, Cat# 15140-122). Human adrenal carcinoma SW-13 cells [ATCC, Cat# CCL-105] were cultured in high-glucose DMEM supplemented with 10% FBS and 1% PS. Human osteosarcoma U2OS cells [ATCC, Cat# HTB-96] were maintained in McCoy’s 5A Medium Modified (Gibco, Cat# 16600082) supplemented with 10% FBS and 1% PS. Human hepatocarcinoma Huh 7.5.1 cells (a generous gift from C. M. Rice, The Rockefeller University, NY, and F. V. Chisari, The Scripps Research Institute, CA) were maintained in high-glucose DMEM (Life Technologies, Cat# 11965-092) supplemented with 10% FBS, 1% PS and 1% non-essential amino acids (Life Technologies, Cat# 11140-050). Human embryonal kidney 293FT cells (ThermoFisher, Cat# R70007) were maintained in high-glucose DMEM supplemented with 10% FBS, 1% PS, and 25 mM HEPES (Life Technologies, Cat# 15630-080). A549 cells [ATCC, Cat# CCL-185] were maintained in DMEM/F-12 (Gibco, Cat# 11320-033) supplemented with 10% FBS (GeminiBio, Cat# 100-106) and 1% PS. Human umbilical vein endothelial cells (HUVEC; Lonza, Cat# C2519A) were maintained in Human Large Vessel Endothelial Cell Basal Medium (Gibco, Cat# M-200-500) supplemented with EGM-SingleQuots (Lonza). Human liver-derived endothelial cells (Lonza, Cat# HLECP1) were maintained in EBM®-2 Endothelial Cell Growth Basal Medium-2 (Lonza, Cat# CC-3156) supplemented with EGM®-2 Endothelial Cell Growth Medium-2 BulletKit (Lonza, Cat# CC-3162) and cultured according to the manufacturer’s protocol. HepaRG cells (ThermoFisher, Cat #HPRGC10) were maintained in Williams E media (ThermoFisher, Cat# 32551020) with HepaRG Growth Medium Supplement (ThermoFisher, Cat# ADD710C). All adherent cells were grown in a humidified incubator at 37°C with 5% CO₂.

### Virus stocks

The authentic CCHFV isolates CCHFV-IbAr10200, CCHFV-Afg09-2990 (labeled as “Afg09”), CCHFV-Turkey2004, CCHFV-Oman-1998091666 (labeled as “Oman”), CCHFV-HY-China, and CCHFV-Kosovo-Hoti (labeled as “Hoti”) were used in this study. Authentic HAZV was acquired from the University of Texas Medical Branch World Reference Center for Emerging Viruses and Arboviruses.

### Authentic HAZV growth, Plaque Assay and Infectivity Measurements

Concentrated stocks of HAZV and CCHFV were grown on SW-13 cells purchased from ATCC (CCL-105). Cells were incubated with the virus for 3 (CCHFV) or 4 (HAZV) days at 37°C in DMEM containing 10% FBS and 1% PS. Following incubation, the HAZV-containing supernatant was centrifuged at 28,000xg for 2 hours at 4°C. Viral pellets were resuspended in NT buffer (10 mM Tris.Cl [pH 7.50], 135 mM NaCl), aliquoted, and stored at -80°C. Following incubation, CCHFV-containing supernatant was centrifuged at 2,000xg for 10 minutes and clarified supernatant was aliquoted and stored at -80°C.

To estimate HAZV virus titer, Vero cells were seeded at a density of 5×10^4^ cells per well in 24-well plates and incubated overnight at 37°C. The next day, HAZV was added at a range of dilutions and incubated at 37°C for 2 hours. Then virus was removed, and the infected cells were washed in phosphate buffered saline (PBS) and overlaid with DMEM containing 1% methylcellulose. Subsequently cells were incubated for 2 days at 37°C and inspected for cytopathic effects. To quantitate infection levels, Vero cells were fixed with 8% paraformaldehyde for 5 minutes, washed and permeabilized with 0.1% Triton X-100 for 15 minutes. The cells were then blocked for 30 minutes with 10% FBS, washed, and stained for 1 hour with HAZV-reactive polyclonal mouse sera obtained from collaborators at the United States Army Medical Research Institute of Allergy and Infectious Disease (USAMRIID). The cells were then washed three times and stained with a fluorescently tagged anti-mouse Alexa Fluor 594 diluted 1:1000 in 10% FBS (ThermoFisher Scientific; A-11005). The cells were then washed three times and stained for cellular nuclei using Hoechst 33342 (ThermoFisher Scientific; 62249). The plates were imaged using a BioTek Cytation 5 cell imager to quantitate focus forming unit titers. Similar cell titer assays were performed for each cell line used in the study. Quantitative infection assays were performed again in a similar manner at an appropriate multiplicity of infection (MOI) that allowed for detection of clear disparate focus forming units.

### Genome-wide CRISPR/Cas9 Knockout Screens

The Brunello CRISPR knockout pooled library in the lentiCRISPR v2 vector was a gift from David Root and John Doench (Addgene, Cat# 73179). SW-13 and U2OS cells expressing the Brunello CRISPR/Cas9 knockout library were generated as previously described [18]. Briefly, the plasmid library was delivered to SW-13 and U2OS cells by lentiviral transduction at a MOI of approximately 0.3 while maintaining 500× library representation with presence of polybrene (7µg/ml; Millipore Sigma, Cat# TR-1003). Two days post-transduction, puromycin hydrochloride (10µg/ml; Mirus Bio, Cat# MIR 5940) was added to the culture medium and maintained until cells in the non-transduced control plate were completely dead (∼48 hours). Library cultures were then expanded until day 12 post-transduction, and SW-13/U2OS–Brunello library cells were cryopreserved at 4 × 10⁷ cells per vial in liquid nitrogen.

Cells were infected with HAZV and recombinant VSV-CCHFV at MOI 0.001 and monitored daily for cytopathic effect. Dead cells were removed; a subset of surviving cells was re-infected to validate resistance to infection. Untreated cells harvested to serve as a reference for sgRNA enrichment analysis. At 21 days post-infection (dpi), surviving cells were pelleted, and genomic DNA (gDNA) was isolated using Blood & Cell Culture DNA Midi Kit (Qiagen, Cat# 13343) according to manufacturer instructions. DNA concentrations were measured with Qubit BR dsDNA quantification kit (Thermo Fisher Scientific, Cat# Q32850) and the QFX fluorometer (DeNovix). For Illumina sequencing, sgRNA sequences were amplified by PCR using Q5 High-Fidelity Mastermix (New England Biolabs, Cat# M0492S) and custom primers binding to conserved sequences flanking the sgRNA in the integrated proviral DNA (Forward: TTGTGGAAAGGACGAAACACCG; Reverse: CCAATTCCCACTCCTTTCAAGACCT). Illumina sequencing adapters (xGen UDI-UMI; Integrated DNA Technologies, Cat# 10005903) were ligated to the purified amplicons using the KAPA HyperPrep Kit (Roche, Cat# KK8502). PCR products were purified and quality-checked using the Bioanalyzer system (Agilent) and subjected to next-generation sequencing (NGS) at the Einstein Genomics Core using an Illumina NextSeq500 DNA sequencer. The resulting FASTQ files were processed with a custom analysis pipeline to extract and quantify sgRNA read counts. Read count files were subsequently analyzed for gene enrichment between CCHFV or HAZV-selected and control populations using the MAGeCK package [27]. Enrichment p-values were generated using GeneRaMeN [28].

### CRISPR/Cas9 engineering of A549, Huh 7.5.1 and Vero knock-out cells

CRISPR sgRNAs targeting AXL and TIM-1 (see Table S1 for sequences) were cloned into the lentiviral vector lentiCRISPR v2 (Addgene, Cat# 52961), which encodes *Streptococcus pyogenes* Cas9. Lentiviral particles were generated in 293FT cells by co-transfecting lentiCRISPR v2 with the packaging plasmid psPAX2 (Addgene, Cat# 12260) and a plasmid encoding vesicular stomatitis virus glycoprotein (VSV-G; Addgene, Cat# 64084). Target cells were then directly exposed to sterile-filtered, lentivirus-laden 293FT cell supernatants in the presence of polybrene (7 μg/ml), and transduced cell populations were selected with puromycin hydrochloride (10 μg/ml). Individual AXL-KO and TIM-1-KO clones were isolated from the respective knockout populations. Genomic DNA was extracted from single-cell clones using the Monarch® Genomic DNA Purification Kit (NEB, Cat# T3010S), following the manufacturer’s instructions. The genomic regions flanking the sgRNA target sites in AXL and TIM-1 were PCR-amplified and sequenced by Oxford Nanopore long read sequencing. Resulting FASTQ files were analyzed to assess frameshift mutations. We identified single-cell clones carrying indels or insertions at the target sites that resulted in frameshifts introducing premature stop codons (Fig. s1).

We confirmed the loss of AXL, TIM-1 protein expression by western blotting and immunofluorescence. Cells were lysed and total protein concentrations were determined using the Pierce™ BCA Protein Assay Kit (Thermo Fisher Scientific, Cat# 23225), with absorbance measured on a Cytation-5 plate reader. Equal amounts of protein were separated by SDS-PAGE on Bolt™ 12% Bis-Tris Plus protein gels (Invitrogen, Cat# NW00120BOX). For western blotting, membranes were probed with the following primary antibodies: LDLR recombinant rabbit monoclonal antibody (Thermo Fisher Scientific, Invitrogen; clone SJ0197, Cat# MA5-32075), goat anti-human AXL (R&D Systems, Cat# AF154), goat anti-human TIM-1 (R&D Systems, Cat# AF1750), and mouse anti-β-actin (Invitrogen, Cat# Ma5-15452). Blots were incubated with HRP-conjugated secondary antibodies specific to rabbit (Santa Cruz Biotechnology, Cat# sc-2357) and goat IgG (Invitrogen, Cat# A15999) and detected using Pierce ECL Plus Western Blotting Substrate (Thermo Scientific, Cat#32132). Signal was visualized using the iBright FL1500 Imaging System (Invitrogen). Knockout efficiency was further validated by immunofluorescence staining using the following polyclonal antibodies: goat anti-human AXL (R&D Systems, Cat# AF154) and goat anti-human TIM-1 (R&D Systems, Cat# AF1750). And the following secondary antibody: Alexa Fluor 488 donkey anti-goat IgG (Invitrogen, Cat# A11055). Representative data are shown in Fig. S1.

### Complementation of A549 Knockout Cell Lines and Generation of AXL/TIM-1 Double Knockout A549 Cells

The full-length AXL coding sequence (CDS) was modified by introducing two-point mutations within the sgRNA target site and PAM sequence to disrupt AXL sgRNA recognition (AXL) was obtained from Twist Bioscience and subcloned into the pLenti-GIII-CMV-Blasti backbone to complement A549 AXL knockout cells described above. Lentivirus was produced by co-transfection of pLenti-GIII-CMV-Blasti-AXL with psPAX2 and VSV-G, as described above. Target cells were transduced with lentiviral supernatant in the presence of polybrene (7 µg/ml) and selected with blasticidin (10 µg/ml). AXL complementation was confirmed with western blotting.

For TIM-1 knockout and AXL-TIM-1 double knockout, A549 and A549 AXL-KO cells were nucleofected with recombinant Cas9 (9 pmol) and sgRNAs (300 pmol each; listed in table s1) using the Lonza Nucleofector Kit T (VCA-1002) on a 2b Nucleofector unit, according to the manufacturer’s instructions. Genetic modification was confirmed by amplicon sequencing (as described above) and Western blotting.

For TIM-1 complementation, full-length TIM-1 CDS (subcloned into the pLenti-GIII-CMV-puromycin backbone) was purchased from Applied Biological Materials Inc. (Cat. no. 23033061). Lentivirus was generated as described above and cells were transduced with the lentiviral supernatant. TIM-1 complementation was confirmed with western blotting.

### Generation of AXL, TIM-1 overexpression cell lines

SW-13 cells, which express low levels endogenous AXL and TIM-1 expression, were used for overexpression studies. The full-length AXL CDS subcloned into the pLenti-GIII-CMV backbone (Applied Biological Materials Inc., Cat. No. 12899061), TIM-1 WT (mentioned above) and TIM-1 N115D/N116D were used to generate lentiviral supernatants as described above and SW-13 cells were transduced with the lentiviral supernatant. Overexpression efficiency was validated with western blotting using the following polyclonal antibodies: goat anti-human AXL (R&D Systems, Cat# AF154) and goat anti-human TIM-1 as described previously.

### Short hairpin RNA (shRNA) mediated gene knockdown on A549, Huh 7.5.1, HUVEC and primary human liver cells

Short-hairpin sequences targeting hAXL and hTIM-1, (subcloned into the pLKO.1 backbone) were purchased from Sigma Millipore (Cat# TRCN0000001040, TRCN0000303580). Scramble shRNA (SCR) was a gift from David Sabatini (Addgene plasmid #1864). 293FT cells were triple transfected with the shRNA plasmids, the lentiviral packaging plasmid psPAX2 (Addgene, Cat#12260), and a plasmid encoding the vesicular stomatitis virus (VSV) G envelope protein (Addgene, Cat#64084). A549, HUVEC, and primary human liver cells were then directly exposed to sterile-filtered, lentivirus-laden 293FT cell supernatants in the presence of polybrene (7 μg/ml), and transduced cell populations were selected with puromycin (10 μg/ml). shRNA knockdown efficiency was first evaluated in A549 cells. After selection, protein expression was assessed by western blotting in A549 and HUVEC cells, while gene expression in primary human liver cells was evaluated by real-time PCR using TaqMan expression assays (Thermo Scientific: Hs01064444_m1 [AXL], Hs00930379_g1 [HAVCR1], and Hg05064520_gH [GAPDH]). Scramble shRNA-transduced cells served as controls for downstream expression analyses. Gene expression changes were quantified using the comparative Ct (ΔΔCt) method, normalized to GAPDH.

### rVSV-CCHFV and tecVLP Infection of cell lines

rVSV-CCHFV and tecVLP pseudoviral systems were utilized to assess infection levels in cell lines and primary cells. rVSV-CCHFV expresses the IbAr10200 CCHFV M-Segment (NC_005300) [25, 26, 33]. tecVLP studies were performed utilizing M segment sequences of IbAr10200, Oman, Kosova-Hoti, and Turkey2004 strain GPCs from GenBank M-segment sequences with accession numbers NC_005300, KR864901, EU037902, and KR864902.1 respectively. These M segments were inserted into the pCAGGS-expression vector and used to generate tecVLPs as described by Zivcec et al [33].

Twenty-four hours prior to infection, cells were seeded in 96 wells (Corning, Cat #3798) at a density of 16,000 cells per well. On the day of infection, rVSV-CCHFV or VLPs were added at their respective MOIs. Plates were incubated for 1 hour in a humidified incubator at 37°C with 5% CO₂.

For rVSV-CCHFV, the viral media was replaced with complete growth medium supplemented with 20 mM ammonium chloride. At 12-16 hours post-infection, cells were fixed with 8% paraformaldehyde in PBS for 5 minutes at room temperature, then washed once with PBS. Cells were permeabilized with 0.1% Triton X-100 in PBS for 10 minutes at room temperature, followed by an additional PBS wash. Hoechst 33342 (ThermoFisher, Cat 62249) was used for nuclear staining and infection was quantified using a Cytation-5 imaging system (Biotek, CYT5FVSN) and Gen5 software by measuring reporter expression.

For VLPs, the viral media is removed, and cells were washed once with 1X PBS. Luciferase activity was measured using Nano-glo luciferase system (Cat #N1110) and read using a Cytation imaging system.

### Authentic CCHFV infection of KO cell lines

Approximately twenty-four hours prior to infection, cells were seeded in 96-well plates (Corning, Cat #3798) at a density of 25,000 cells per well. On the day of infection, authentic CCHFV (strains IbAr10200, Afg09, HY-China, Turkey, Oman, and Kosova-Hoti) were added at their respective MOIs and incubated at 37°C. After 48 hours (strains IbAr10200, Afg09, HY-China) or 72 hours (strains Turkey, Oman, Kosova-Hoti) post-infection, the culture medium was removed, and plates were submerged in 10% neutral buffered formalin and fixed for at least 24 hours at room temperature. Plates were removed from formalin, washed with PBS three times, and permeabilized with 0.2% Triton-X for 10 minutes at room temperature. Following permeabilization, plates were washed three times with PBS and blocked with blocking buffer (Cell Stain Buffer; ThermoFisher Scientific) for 2 hours at 37°C. Infected cells were detected by successive incubation with CCHFV-specific anti-nucleocapsid antibody 9D5 (3 µg/mL; BEI NR-40270) and secondary detection antibody (goat anti-mouse) conjugated to AlexaFluor 488 (diluted 1:2,000; Invitrogen). Percent infection was determined using a Cytation 5 high-content imaging instrument and data analysis was performed using Gen5.11 software (BioTek).

### PtdSer and PtdCho liposome blocking

Twenty-four hours prior to infection Vero (rVSV-CCHFV) or VeroE6 (authentic CCHFV) cells were seeded in 96 wells (Corning, Cat #3798) at a density of 16,000 or 25,000 cells per well, respectively. On the day of infection cells were pretreated with liposomes composed of phosphatidyl-serine (PtdSer) or phosphatidyl-choline (PtdCho). PtdSer and PtdCho liposomes were generated as previously described (10.1016/j.immuni.2007.11.011). Liposomes were added in a dilution series starting at 100 uM diluted 2-fold down the plate, cells were incubated with liposomes for 1 hour at 37°C5% CO₂. Following incubation rVSV-CCHFV was added to cells at MOI 10 or authentic CCHFV strains IbAr10200, Afg09, Turkey2004, Oman and HY-China at MOI 0.08, 0.2, 0.02, 0.2, or 0.1, respectively and incubated for 1 hour at 37°Cat 5% CO₂. For rVSV-CCHFV, the viral media was replaced with complete growth medium supplemented with 20 mM ammonium chloride. At 12-16 hours post-infection, cells were fixed with 8% paraformaldehyde in PBS for 5 minutes at room temperature, then washed once with PBS. Cells were permeabilized with 0.1% Triton X-100 in PBS for 10 minutes at room temperature, followed by an additional PBS wash. Hoechst 33342 (ThermoFisher, Cat 62249) was used for nuclear staining, and infection was quantified using a Cytation imaging system (Agilent, CYT5FVSN) by measuring reporter expression. For authentic CCHFV, the viral media was replaced with complete growth media. After 48 hours (strains IbAr10200, Afg09, HY-China) or 72 hours (strains Turkey2004, Oman, Kosova-Hoti), the culture medium was removed, and plates were submerged in 10% neutral buffered formalin and fixed for at least 24 hours at room temperature. Plates were removed from formalin, washed with PBS three times, and permeabilized with 0.2% Triton-X for 10 minutes at room temperature. Following permeabilization, plates were washed three times with PBS and blocked with blocking buffer (Cell Stain Buffer; ThermoFisher Scientific) for 2 hours at 37°C. Infected cells were detected by successive incubation with CCHFV-specific anti-nucleocapsid antibody 9D5 (3 µg/mL; BEI NR-40270) and secondary detection antibody (goat anti-mouse) conjugated to AlexaFluor 488 (diluted 1:2,000; Invitrogen). Percent infection was determined using a Cytation imaging system and data analysis was performed using Gen5.11 software (Agilent).

### Antibody blocking

Twenty-four hours prior to infection cells were seeded in 96 wells (Corning, Cat #3798) at a density of 16,000 cells per well. On the day of infection cells were pretreated with polyclonal antibodies targeting AXL (R&D Systems, AF154) and TIM-1 (R&D Systems, AF1750).

Antibodies were added in a dilution series starting at 10ug/mL and diluted 2-fold down the plate, cells were incubated with liposomes for 1 hour at 37 degrees 5% CO₂. Following incubation rVSV-CCHFV was added to cells at MOI 10 and incubated for 1 hour at 37 degrees at 5% CO₂. For rVSV-CCHFV, the viral media was replaced with complete growth medium supplemented with 20 mM ammonium chloride. At 12-16 hours post-infection, cells were fixed with 8% paraformaldehyde in PBS for 5 minutes at room temperature, then washed once with PBS. Cells were permeabilized with 0.1% Triton X-100 in PBS for 10 minutes at room temperature, followed by an additional PBS wash. Hoechst 33342 (ThermoFisher, Cat 62249) was used for nuclear staining and infection was quantified using a Cytation imaging system (Biotek, CYT5FVSN) by measuring reporter expression.

### Bemcentinib blocking

Twenty-four hours prior to infection cells were seeded in 96 wells (Corning, Cat #3798) at a density of 16,000 cells per well. On the day of infection cells were pretreated with bemcentinib. Bemcentinib was added in a dilution series starting at 10ug/mL and diluted 2-fold down the plate, cells were incubated with liposomes for 1 hour at 37 degrees 5% CO₂. Following incubation rVSV-CCHFV was added to cells at MOI 10 and incubated for 1 hour at 37 degrees at 5% CO₂. For rVSV-CCHFV, the viral media was replaced with complete growth medium supplemented with 20 mM ammonium chloride. At 12-16 hours post-infection, cells were fixed with 8% paraformaldehyde in PBS for 5 minutes at room temperature, then washed once with PBS. Cells were permeabilized with 0.1% Triton X-100 in PBS for 10 minutes at room temperature, followed by an additional PBS wash. Hoechst 33342 (ThermoFisher, Cat 62249) was used for nuclear staining, and infection was quantified using a Cytation imaging system (Biotek, CYT5FVSN) by measuring reporter expression.

### Mouse models

#### Ethics

Animals had previously never undergone experimentation. Animals were provided with food and water ad libitum and housed in individually ventilated cages under specific pathogen-free conditions at USAMRIID. Murine challenge studies were conducted under an Institutional Animal Care and Use Committee (IACUC)-approved protocol in compliance with the Animal Welfare Act, Public Health Service Policy, and other federal statutes and regulations relating to animals and experiments involving animals. The facility where this research was conducted (USAMRIID) is accredited by AAALAC International and adheres to principles stated in the Guide for the Care and Use of Laboratory Animals, National Research Council 2011.

Animal research and breeding performed at the University of Iowa (UI) were conducted in accordance with the Animal Welfare Act and the recommendations in the Guide for the Care and Use of Laboratory Animals of the National Institutes of Health (PHS Animal Welfare Assurance Number: D16-00009, A3021-01) and with the IACUC guidelines (protocol #: 4021280, Filovirus glycoprotein/cellular protein interactions and virus glycoprotein/cellular protein interactions).

### Generation and breeding of murine lines

WT C57BL/6 mice were purchased from Jackson Lab (Bar Harbor, ME). C57BL/6J *Ifnar^−/–^* mice were a kind gift from Dr. John Harty (University of Iowa, Iowa City, IA, USA). All C57BL/6 *Ifnar^−/–^* lines were bred at the University of Iowa. C57BL/6 *Ifnar^−/–^*, *Havcr1* (*TIM-1)^− /–^ Ifnar^−/–^*, and *Timd4* (*TIM-4)^−/–^ Ifnar^−/–^* have been previously described (PMID: 31825972, PMID: 31242184, PMID: 20368430, PMID: 22144095). C57BL/6 *AXL^−/–^* were obtained from Dr. Rolf Brekken (Univ. Texas, Southwestern, Houston, TX) and crossed onto a C57BL/6 *Ifnar^−/–^* background, generating C57BL/6 *AXL^−/–^ Ifnar^−/–^* (PMID: 33811159, PMID: 40938099). Triple knockout C57BL/6 *TIM-1^−/–^ AXL^−/–^ Ifnar^−/–^* (*TIM-1^−/–^* | *AXL^−/–^*) were generated by crossing C57BL/6 *TIM-1^−/–^ Ifnar^−/–^* with C57BL/6 *AXL^−/–^ Ifnar^−/–^* (PMID: 40938099). Triple knockout C57BL/6 *TIM-4^−/–^ AXL^−/–^ Ifnar^−/–^* (*TIM-4^−/–^*| *AXL^−/–^*) were generated by crossing C57BL/6 *TIM-4^−/–^ Ifnar^−/–^* with C57BL/6 *AXL^−/–^ Ifnar^−/–^*. Mouse genotypes were verified using primers described in Table S1D and PCR conditions previously described (PMID: 31242184). Briefly, as approved and described by the UI IACUC policies and guidelines, DNA is extracted from a small (<5 mm) portion of the end of the mouse tail. Tissue was processed using the Wizard Genomic DNA Purification Kit by Promega as described by the manufacturer’s instructions. DNA is quantified by nanodrop. For each reaction, 100 ng of genomic DNA, primers, and the New England Biolabs OneTaq 2X Master Mix are mixed, and a touchdown PCR protocol is used as recommended by The Jackson Laboratory.

### Animal challenge

8-24-week-old male and female IFNAR^-/-^, IFNAR^-/-^/AXL^-/-^, IFNAR^-/-^/TIM-1^-/-^, IFNAR^-/-^/TIM-4^-/-^, IFNAR^-/-^/AXL^-/-^/TIM-1^-/-^, and IFNAR^-/-^/AXL^-/-^/TIM-4^-/-^ mice were exposed intraperitoneally (IP) to 10 plaque forming units (PFU) of CCHFV-Turkey2004. At 5 days post-challenge, n=5 mice per strain were euthanized and whole blood, the liver, the spleen, and the kidneys were collected. Mice were sampled at random. The remaining animals were observed daily for clinical signs of disease and morbidity for 28 days post-challenge. Mice were scored on a 4-point IACUC-approved grading scale: 1 is decreased grooming and/or ruffled fur; 2 is subdued behavior when un-stimulated, 3 is lethargy, hunched posture, and/or subdued behavior even when stimulated, and 4 is bleeding, unresponsiveness, severe weakness, and/or inability to walk. Mice scoring a 4 were considered moribund and euthanized based on IACUC-approved methods. Mice scoring a 3 were observed at a minimum of twice daily. The animal study protocol was approved by the Institutional Review Board of the USAMRIID Institutional Animal Care Committee (protocol code AP-25-001, date of approval 15 January 2025).

### Plaque assay

Whole blood was centrifuged at 12,000 x g for 10 minutes to collect serum. Tissues were homogenized in infection media in gentleMACS M tubes (Miltenyi Biotec) to generate a 10% tissue homogenate and clarified to remove cellular debris by centrifugation at 4,000 x g for 10 minutes. 10-fold serial dilutions of serum and tissue homogenates were prepared in infection media and 200 µL of each dilution was inoculated onto SW-13 cell monolayers in 6-well plates. After adsorption for 1 hour at 37°C, cell monolayers were overlaid with a mixture of 1 part 1% agarose (Seakem ME) and 1-part 2x Eagle basal medium, 30 mM HEPES buffer, and 2% heat-inactivated FBS. Plates were incubated for 4 days at 37°C, at which point a second overlay supplemented with 5% neutral red was added. Plaques formed by infectious CCHFV were counted 8 hours later and titers are shown as PFU/mL.

### PCR

Tissue samples were homogenized and clarified prior to extraction as required. Samples were inactivated using a 3:1 ratio of TRIzol LS Reagent to sample (ThermoFisher, Waltham, MA). 400 µL of Trizoled material was purified using the EZ1 Virus Mini Kit v 2.0 (Qiagen) using the EZ1 Advanced XL robot (Qiagen) according to the manufacturer’s recommendations. The total elution volume was 150 µL. RT-qPCR was performed using TaqPath™ 1-Step RT-qPCR Master Mix, CG (ThermoFisher) on the QuantStudio DX (ThermoFisher) with primers and probes targeting the L segment of CCHFV (Forward – 5’-CCCTTTGGTAGGCTGTTT-3’, Reverse – 5’-CTGTTCATGTGAAGTGTTTGAG-3’, Probe – 6FAM 5’-AGCTCGGCGGTTGCCTTGGAAGATGCTGAA-3’ TAMRA). Reference material was quantified using the QIAcuity Digital PCR System and the QIAcuity One-Step Viral RT-PCR Kit (Qiagen) using the same RT-qPCR assay. Samples were run in technical triplicate and concentrations were calculated using a standard curve of quantified reference material included on each run. Sample concentrations are reported in copies of target per ml and are based on extracted material provided within the standard curve. Concentrations that fell outside the linear range of the standard curve were extrapolated. The limit of detection of the assay based on the standard curve is approximately 5 copies/µl.

### Pathology

We performed pathological analysis on animals sacrificed 5 days post-infection. Tissues were trimmed, processed, embedded in paraffin, sectioned at 5 µm, stained with hematoxylin and eosin (H&E), coverslipped, and evaluated for histopathologic lesions. Replicate sections were placed on positively charged slides and stained by RNA *in situ* hybridization (ISH). Detection of viral RNA was performed using the RNAscope 2.5 High Definition (HD) RED Assay (Advanced Cell Diagnostics, Newark, CA, USA). Briefly, a custom ISH target probe (Advanced Cell Diagnostics, Cat No. 1907351-C1) was designed and synthesized to target the 489–1995 nucleotide region of the Crimean-Congo hemorrhagic fever virus (CCHFV) Turkey2004 isolate genome (GenBank accession no.KR864902.1). Tissue sections were deparaffinized with xylene, underwent a series of ethanol washes and peroxidase blocking, and were then heated in kit-provided antigen retrieval buffer and then digested by kit-provided proteinase. Sections were then incubated with the target probe at 40°C in a hybridization oven for 2 h. After rinsing, ISH signal was amplified using kit-provided Pre-amplifier and Amplifier conjugated to alkaline phosphatase and incubated with a Fast Red substrate solution (Abcam, Cat ab64254) for 10 min at room temperature. Sections were then stained with hematoxylin, air-dried, and coverslipped. All slides were examined by a board-certified veterinary pathologist.

### Statistical Analyses

Differences among multiple groups were assessed using the Kruskal–Wallis test for nonparametric data or analysis of variance (ANOVA) for parametric data. Following ANOVA, Dunnett’s test was used to compare each treatment group with the control group, and Tukey’s test was used for all pairwise group comparisons. Comparisons between two independent groups with unequal variances were performed using Welch’s t-test. Statistical significance was defined as *p*≤ 0.05. Survival outcomes were compared against each group by Mantel-Cox tests. Weight loss and clinical outcomes were compared against each group by the Analysis of Variance (ANOVA) test. Statistical significance was determined by p ≤ 0.05, *, p < 0.05, **, p < 0.01, ***, P<0.001 and ****, p<0.0001. All statistical analyses were conducted in the Prism Software V9.5.1 (GraphPad).

### Biosafety considerations

All authentic virus manipulation and *in vivo* experimentation with CCHFV were conducted in a biosafety level 4 (BSL-4) environment following all institutional biosafety Standard Operating Procedures (SOPs) and regulatory requirements and in compliance with the CDCs Biosafety in Microbiological and Biomedical Laboratories (BMBL) guidelines. Animal experiments were conducted in the ABSL-4 facility and were approved by USAMRIID IACUC. Inactivated samples were transferred from the BSL-4 facility to BSL-2 laboratories following validated protocols approved by the Institutional Biosafety Committee at USAMRIID.

## Author Contributions

Conceptualization: J.B., S.R.M., W.J.M., A.S.H., K.C., E.K.

Investigation: J.B., S.R.M., T.P., A.I.K., R.R.B., M.D.R., P.T.D., C.P.S., S.B.S., X.Z., S.R.C., K.C., E.K.

Methodology: J.B., S.R.M., M.D.R., P.T.D., C.P.S., X.Z., W.M., K.C., E.K.

Supervision: S.R.M.,W.J.M., C.P.S., X.Z., A.S.H., K.C., E.K.

Visualization: J.B., S.R.M., P.T.D., C.P.S., S.B.S., X.Z., W.J.M., K.C., E.K.

Writing – original draft: J.B., S.R.M., K.C., E.K. Writing – editing: all authors

## Acknowledgements

We thank Matt Ramirez, Narisa Lee, and Karyme Paez (Einstein), and Cecilia O’Brien and Kandis Cogliano (USAMRIID) for laboratory management and administrative support. We also thank the following divisions at USAMRIID: Veterinary Medicine Division and Statistics Division. We thank Ms. Holly Bloomfield, Mr. Neil Davis, and Ms. Ashley Bryan (United States Army Medical Research Institute of Infectious Diseases [USAMRIID], Fort Detrick, Frederick, MD, USA) for their histologic or molecular pathology support. The opinions, interpretations, conclusions, and recommendations presented are those of the author and are not necessarily endorsed by the U.S. Army or Department of Defense. This work was partially supported by U.S. National Institute of Health grants U19AI181977 (to K.C., S.R.M., and A.S.H.). J.B. was additionally supported by the NIH training grant T32AI070117-17 at Albert Einstein College of Medicine. The opinions, interpretations, conclusions, and recommendations presented are those of the authors and are not necessarily endorsed by the U.S. Army, Department of Defense, or the US Department of Health and Human Services. The use of either trade or manufacturers’ names in this report does not constitute an official endorsement of any commercial products. This report may not be cited for purposes of advertisement. This does not constitute an endorsement by the U.S. Government of this or any other contractor.

