## Supplementary Figures for "Phosphatidylserine receptors are critical for lethal infection by Crimean-Congo hemorrhagic fever virus"

Fig S1

A549 WT : CATCCTCTTTCTCCTGCGAAGCCCATAACGCCAAGGGGGTCACCACATCCCGCACAGCCAC
AXL KO : CATCCTCTTTCTCCTGCGAAGCCCATAACGC-----Δ16-----TCCCGCACAGCCAC
: CATCCTCTTTCTCCTGCGAAGCCCATAACGC- AAGGGGGTCACCACATCCCGCACAGCCAC

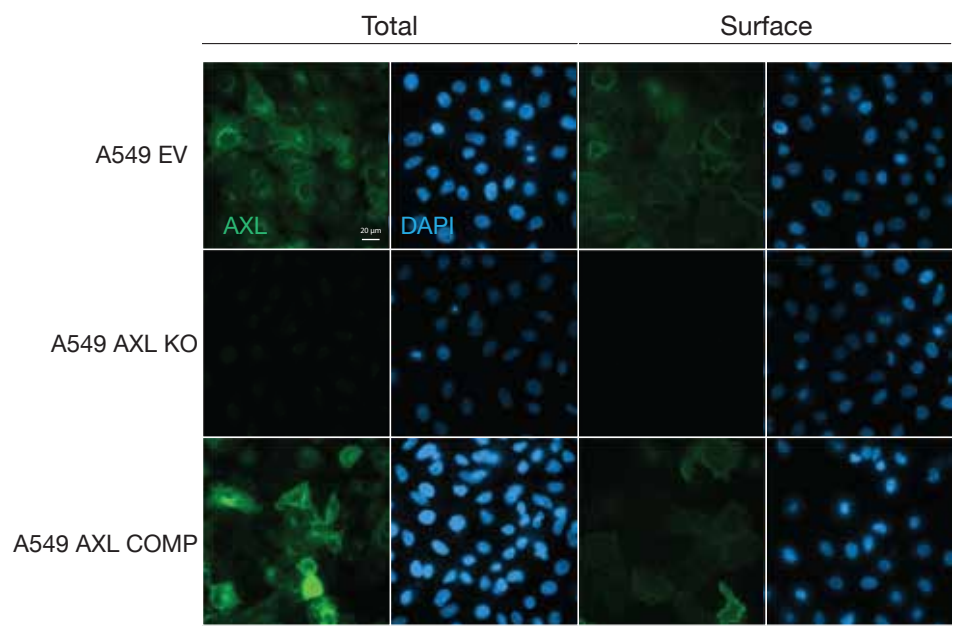

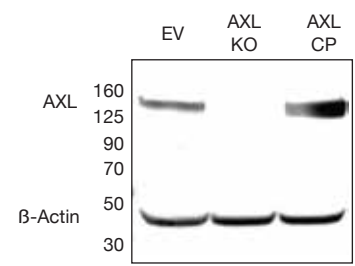

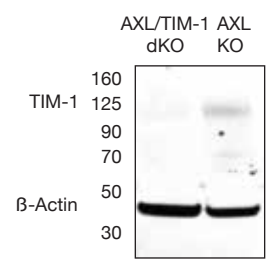

A549 WT : GGACACACGCTATAAGCTATTGGGGGACC.....GCACCGTGGGTGGTTCAATGACATGA
TIM-1 KO : GGACACACGCTATAAGCTATT-----Δ93-----GTGGTTCAATGACATGA

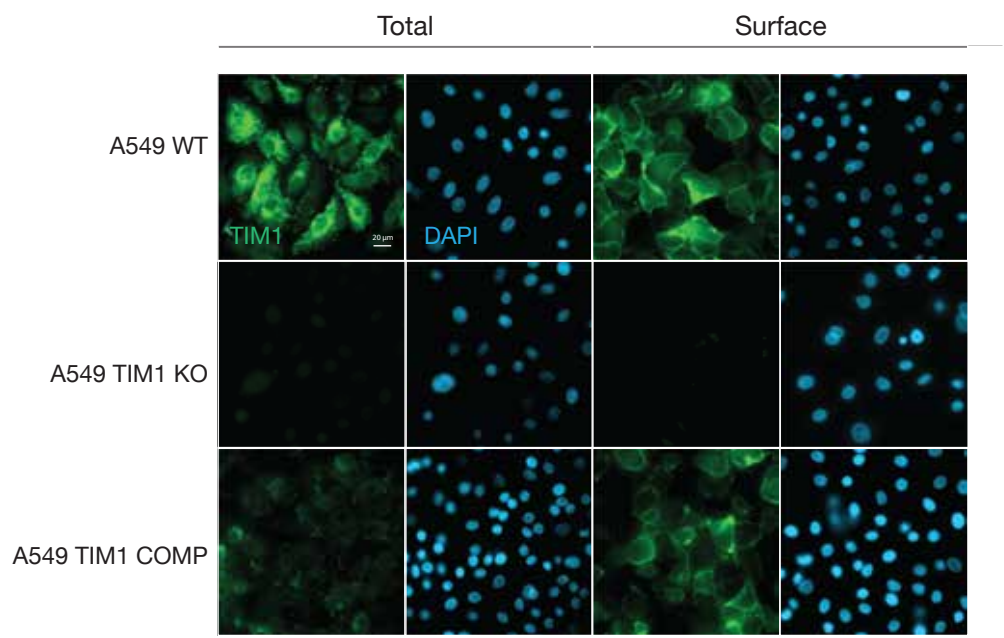

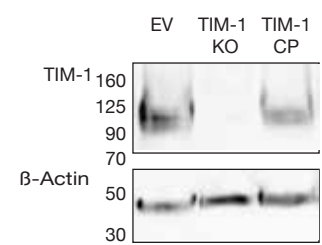

### **Supplementary Figure 1: Confirmation of AXL and TIM-1 knockout and complementation in A549 cells**

Single cell CRISPR/Cas9 knockout and complement A549 cells were sequenced via Oxford nanopore long read sequencing for knockout at specific gRNA locus. Cells were stained via immunofluorescent microscopy for AXL/-TIM-1 expression levels. Cells were tested for AXL/TIM-1 expression levels via western blot.

Vero WT : CTCTTGCCCTGTCAGAATCGCCTCCCTACAGCTTTCCGATGCGGGACAGTACCAGTGTTT
AXL KO : CTCTT-----Δ29-----TCCGATGCGGGACAGTACCAGTGTTT
: CTCTTGCCCTGTCAGAATCGCCCAATTAGGGAA---Δ7---AGCTTTCCGATGCGGGACAGTACCAGTGTTT
Total Surface

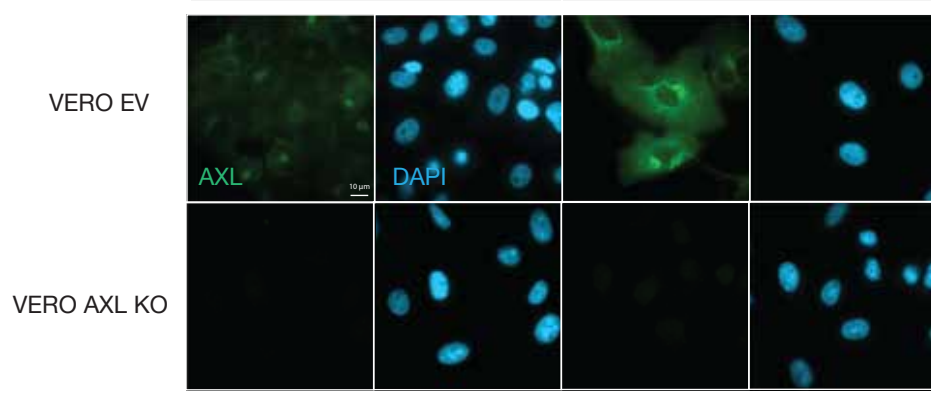

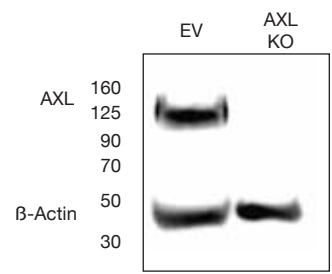

Vero WT : ACTCCAATTGTCAGAACTGTTCGAA.....CCAACGACAACCATGGTTTCTACCTTT
TIM-1 KO : ACTCCAATT-----Δ465-----ACAACCATGGTTTTCTACCTTT
: ACTC- - - TGT- - GAACTGTTTCGAA.....AACGGACAACCATGGTTTTCTACCTTT

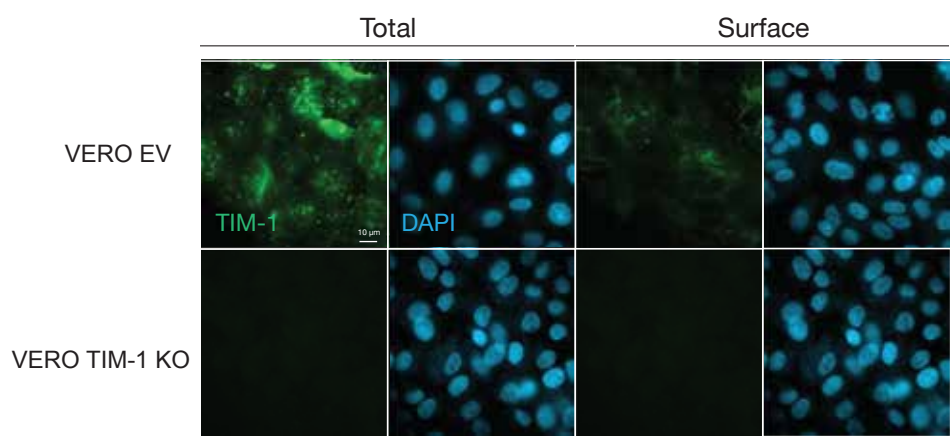

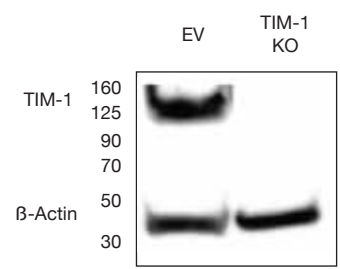

Huh 7.5.1 WT : GGACACACGCTATAAGCTATTGGGGGACC.....GCACCGTGGGTGGTTCAATGACATGA
TIM-1 KO : GGACACACGCT-----Δ20-----GCACCGTGGGTGGTTCAATGACATGA
: GGACACACGCTATAAGCTATTGGGGGACC.....GCACCGTGGGTGGTTCAATGACATGA

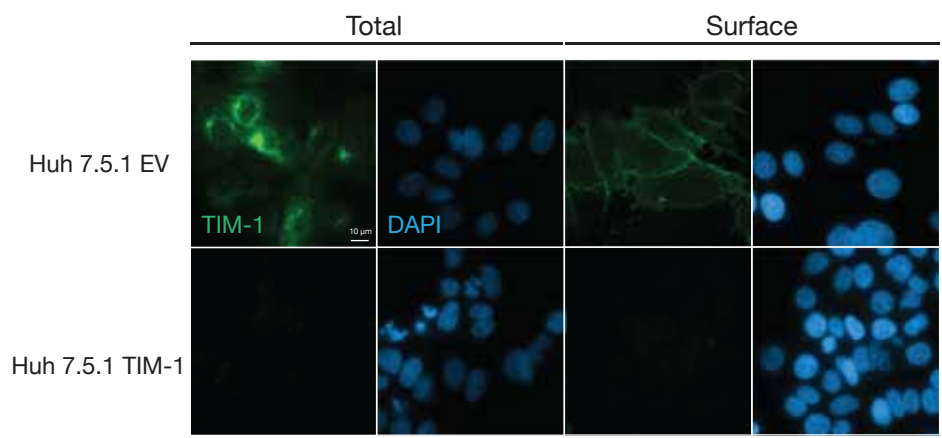

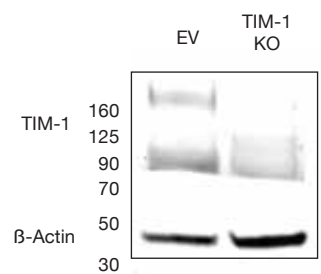

**Supplementary Figure 2: Confirmation of AXL and TIM-1 knockout in Vero cells and confirmation of TIM-1 knockout in HUH 7.5.1 cells.**

Single cell CRISPR/Cas9 knockout Vero cells were sequenced via Oxford nanopore long read sequencing for knockout at specific gRNA locus. Cells were stained by immunofluorescent microscopy for AXL/TIM-1 expression levels. Cells were tested for AXL/TIM-1 expression levels via western blot. HUH 7.5.1 cells were analyzed via sequencing, immunofluorescent staining and western blot for TIM-1 expression levels.

Fig S3

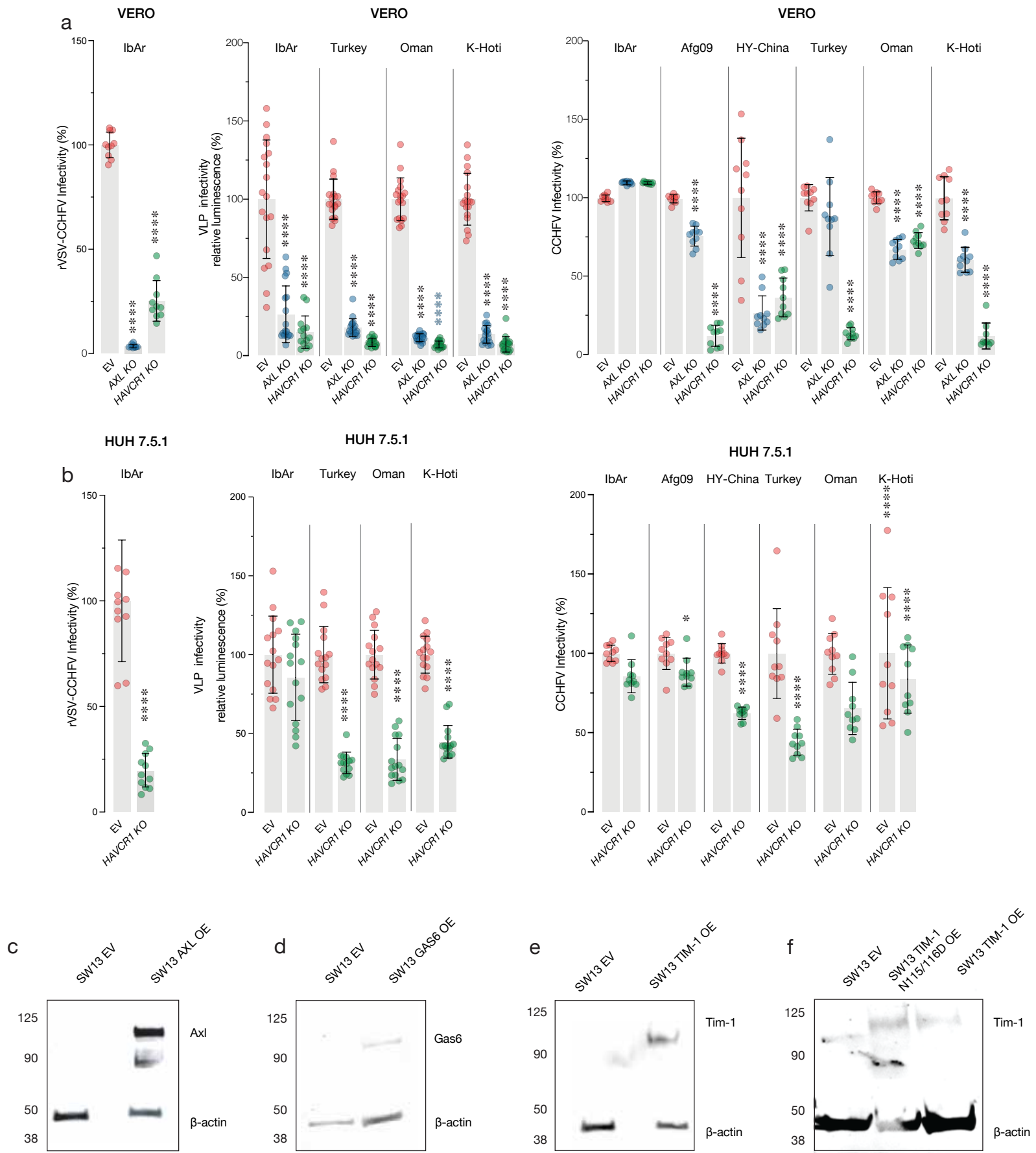

**Supplementary Figure 3: Infection of Vero/HUH7.5.1 knockout cells and confirmation of overexpression in SW-13 cells.**

Infection of Vero cell clones knocked out (KO) for *AXL* or *HAVCR1* (encoding TIM-1) and HUH7.5.1 cells KO for *HAVCR1*. **(a,b left panel)** Cells were exposed to rVSV-CCHFV GPC (multiplicity of infection, MOI of 0.2 infectious units [IU]/cell) for 1 h prior to addition of $\text{NH}_4\text{Cl}$ to stop infection. Infected cells were enumerated at 16 hpi by immunofluorescence microscopy. **(a,b middle panel)** Cells were exposed to equal volumes of $\text{TCID}_{50}$-normalized CCHFV tecVLPs, and transduction was measured at 16 hpi by luminometry. **(a,b right panel)** Cells were exposed to authentic CCHFV (MOI, 0.2 PFU/cell), and infected cells were enumerated by immunofluorescence microscopy. Means ± s.d. from three independent experiments are shown. Groups (WT vs. others) were compared by one-way ANOVA with Dunnett's post hoc test. Groups (Huh 7.5.1 vs TIM-1 KO) were compared by unpaired t-test with Welch's correction. SW-13 cells were probed for overexpression of **(c)** AXL, **(d)** Gas6, **(e)** TIM-1 and **(f)** TIM-1 N115/116D via western blot analysis.

Fig S4

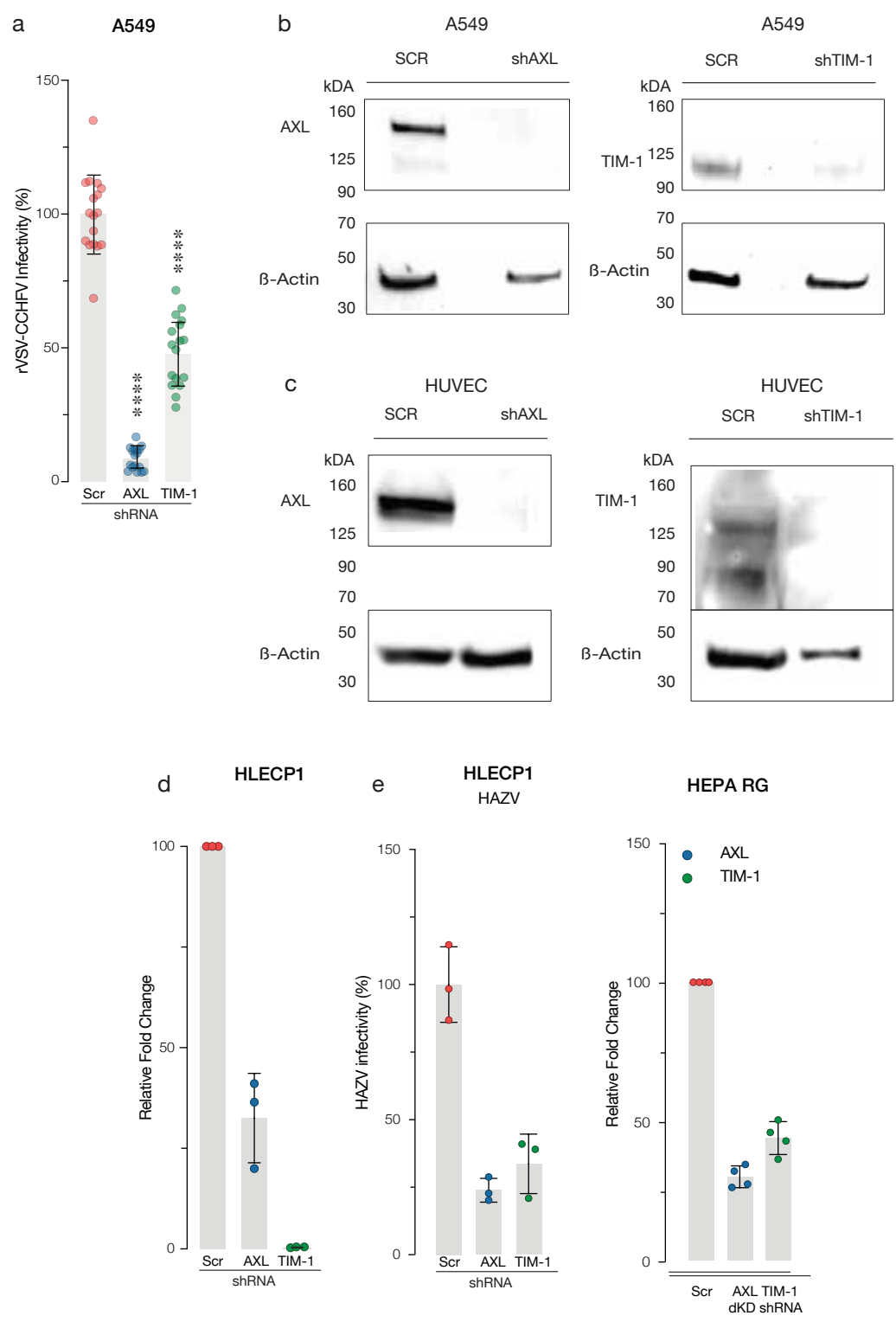

**Supplementary Figure 4: Confirmation of short hairpin knockdown in A549 and HUVEC cells, Confirmation of Knockdown in HLECP1 cells and HAZV infection.**

Infection of A549 cells with shRNA knockdown. **(a)** Cells were exposed to rVSV-CCHFV GPC (multiplicity of infection, MOI of 0.2 infectious units [IU]/cell) for 1 h prior to addition of NH<sub>4</sub>Cl to stop infection. Infected cells were enumerated at 16 h hpi by immunofluorescence microscopy. **(b,c)** Western blot confirmation of AXL and TIM-1 protein expression in WT cells and shRNA knockdown cells. **(d)** qPCR confirmation of HLECP1 knockdown of AXL and TIM-1. **(e)** cells were exposed to HAZV (multiplicity of infection, MOI of infection 1) for 1 hour prior to addition of NH<sub>4</sub>Cl to stop infection. Infected cells were enumerated at 16 hpi by immunofluorescence microscopy. Data, (by two-way ANOVA with Tukey's post hoc test. **(f)** qPCR confirmation of HepaRG knockdown of AXL and TIM-1.

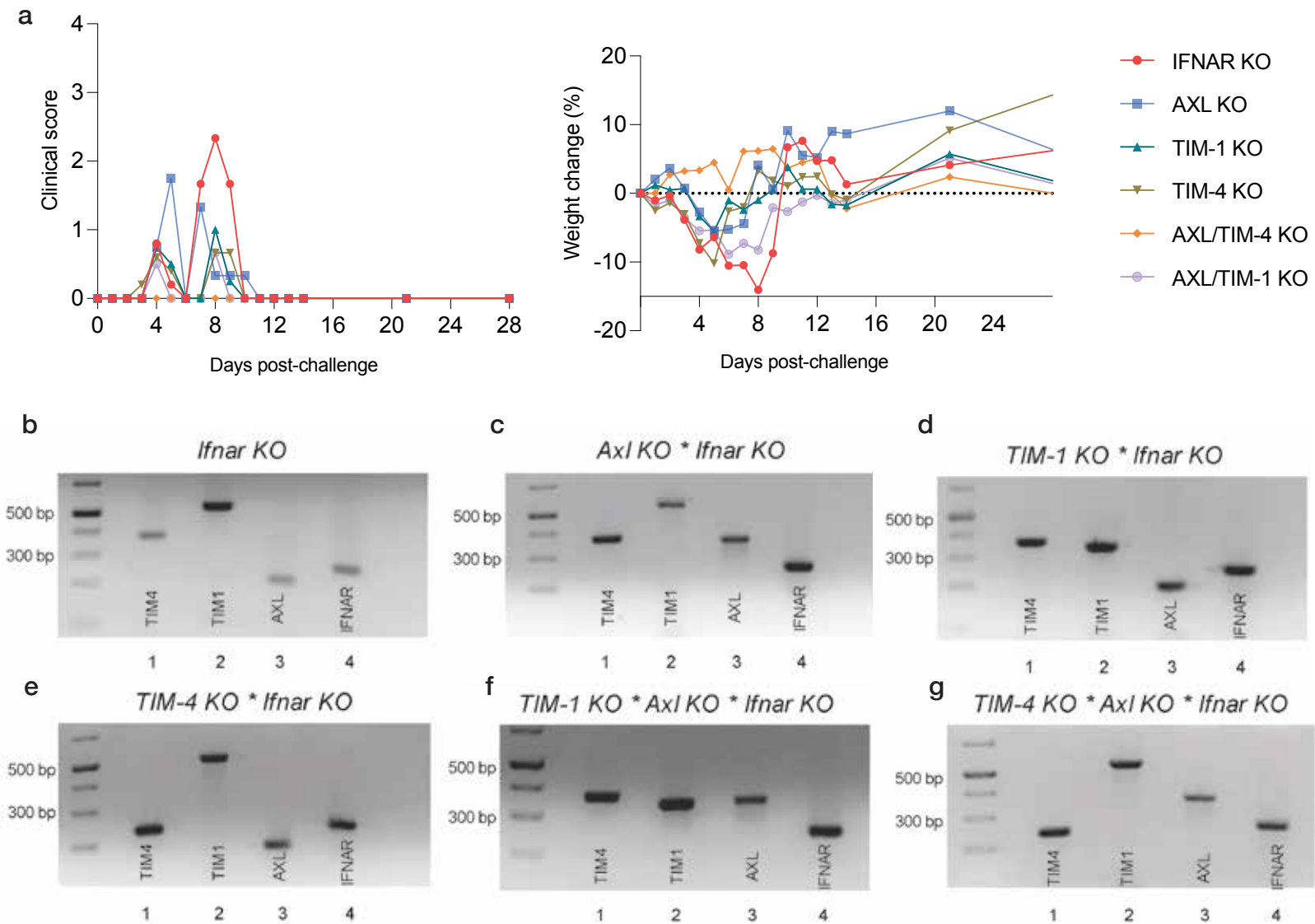

### Supplementary Figure 5: Clinical Scoring, Weight loss and Genotyping analysis of PS receptor deficient mice.

IFNAR (-/-) background wildtype and knockout mice (n=15-16 per strain) were inoculated with 10 PFU for Turkey2004 via the i.p. route. 5 days post-challenge, n=5 mice per strain were euthanized and harvested for serum, liver, kidney, and spleen. Mice were monitored for (a) clinical signs of disease, and (b) weight loss for 28 days post-challenge. Representative images of amplified mouse genomic DNA for: *Ifnar KO* (c), *Axl KO \* Ifnar KO* (d), *Havcr1 KO \* Ifnar KO* (e), *Timd4 KO \* Ifnar KO* (f), *Havcr1 KO \* Axl KO \* Ifnar KO* (g), and *Timd4 KO \* Axl KO \* Ifnar KO* (h). The mutant *Timd4* gene fragment (lane 1) is 250 base pair (bp) and wild type is 373 bp. The mutant *Havcr1* gene fragment (lane 2) is 330 bp and wild type is 536 bp. The mutant *Axl* gene fragment (lane 3) is 333 bp and wild type is 196 bp. The *Ifnar* mutant gene fragment (lane 4) is 250 bp and wild type is 155 bp.
